# Ribosome rescue inhibitors clear *Neisseria gonorrhoeae in vivo* using a new mechanism

**DOI:** 10.1101/2020.06.04.132530

**Authors:** Zachary D. Aron, Atousa Mehrani, Eric D. Hoffer, Kristie L. Connolly, Matthew C. Torhan, John N. Alumasa, Pooja Srinivas, Mynthia Cabrera, Divya Hosangadi, Jay S. Barbor, Steven C. Cardinale, Steven M. Kwasny, Lucas R. Morin, Michelle M. Butler, Timothy J. Opperman, Terry L. Bowlin, Ann Jerse, Scott M. Stagg, Christine M. Dunham, Kenneth C. Keiler

## Abstract

The *trans*-translation pathway for rescuing stalled ribosomes is conserved and essential in bacterial pathogens but has no mammalian homolog, making it an ideal target for new antibiotics. We previously reported the discovery of a family of acylaminooxadiazoles that selectively inhibit *trans*-translation, resulting in broad-spectrum antibiotic activity. Optimization of the pharmacokinetic and antibiotic properties of the acylaminooxadiazoles produced MBX-4132, which cleared multiple-drug resistant *Neisseria gonorrhoeae* infection in mice after a single oral dose. Cryo-EM studies of non-stop ribosomes showed that acylaminooxadiazoles bind to a unique site near the peptidyl-transfer center and significantly alter the conformation of ribosomal protein L27, suggesting a novel mechanism for specific inhibition of *trans*-translation by these molecules.

**One Sentence Summary:** Ribosome rescue inhibitors reveal a new conformation of the ribosome and kill drug-resistant *Neisseria gonorrhoeae in vivo*.

## Main text

Antibiotic-resistant bacterial pathogens pose a substantial threat to human health and are projected to cause up to 10 million deaths per year by 2050 if new antibiotics are not developed (*1*). Among the 5 most dangerous “Urgent Threats” identified by the CDC is drug-resistant *Neisseria gonorrhoeae*, which infects >500,000 people per year in the US (*2*). Bacterial ribosome rescue pathways have been proposed as potential antibiotic targets because they are essential in bacteria and are highly dissimilar from cytoplasmic mechanisms to disassemble non-stop ribosomes in eukaryotes (*3*). Bacterial ribosomes frequently stall at the 3’ end of mRNAs lacking an in-frame stop codon, due to physical or nucleolytic damage to the mRNA or premature termination of transcription (*4*). These “non-stop” ribosomes must be rescued to maintain protein synthesis capacity and cell viability (*4*). All bacterial species that have been studied use *trans*-translation as the primary ribosome rescue pathway. During *trans*-translation, transfer-messenger RNA (tmRNA) and the protein SmpB recognize non-stop ribosomes and use tRNA-like and mRNA-like properties of tmRNA to add a short sequence to the nascent polypeptide and terminate translation at a stop codon within tmRNA (*4*). In some bacteria, including *N. gonorrhoeae*, tmRNA and SmpB are essential (*5, 6*). Other species have an alternative ribosome rescue factor (ArfA, ArfB, ArfT, or BrfA) that acts as a backup system for rescuing ribosomes when *trans*-translation activity is not sufficient (*7–10*). Deletions of the alternative ribosome rescue factors and tmRNA or SmpB are synthetically lethal, indicating that at least one mechanism for ribosome rescue is required for bacterial viability (*5*).

A family of acylaminooxadiazoles identified in a high-throughput screen for inhibitors of *trans*-translation in *Escherichia coli* was shown to have broad-spectrum antibiotic activity *in vitro* (*3, 11*). Experiments in *E. coli* and *Mycobacterium smegmatis* showed that KKL-2098, a cross-linkable acylaminooxadiazole derivative, bound 23S rRNA near the peptidyl-transfer center (PTC), suggesting that these molecules inhibit *trans*-translation by binding the ribosome (*11*). Based on these data, we sought to optimize the acylaminooxadiazole activity against *N. gonorrhoeae* and establish the basis for selective inhibition of *trans*-translation.

Evaluation of *in vitro* pharmacokinetic properties of the original hit, KKL-35, revealed that the amide bond is rapidly hydrolyzed in liver microsomes, making it unsuitable for animal studies (Fig. 1, Table S1). To enable animal studies, >500 analogs of KKL-35 were designed and evaluated for potency, toxicity, and pharmacokinetic properties (Fig. 1, Table S1). Compound potency, assessed by minimum inhibitory concentration (MIC) against *N. gonorrhoeae* and activity (EC_50_) in a *trans*-translation luciferase reporter assay (*3*), was responsive to structural changes, consistent with specific binding of the molecule to the target (Fig. 1, Table S1). Conceptually, the compound can be divided into 4 distinct zones (Fig. 1A). The central portion (Zones 2 and 3) played a critical role in activity, and the termini (Zones 1 and 4) tolerated changes that facilitated tuning physical properties and potency. A key finding of these experiments was that replacement of the Zone 3 amide with a urea dramatically improved metabolic stability without significantly decreasing potency (Fig. 1, Table S1). MBX-4132, a uriedooxadiazole, exhibited excellent stability in both murine liver microsomes and murine serum as well as excellent Caco-2 permeability (Fig. 1B, Table S2). MBX-4132 inhibited *trans*-translation both in the *E. coli* luciferase reporter assay (Fig. 1B) and in an *in vitro* reconstituted assay (IC_50_ = 60 nM), but did not inhibit translation *in vitro* (Fig. 1D).

**Figure 1.**
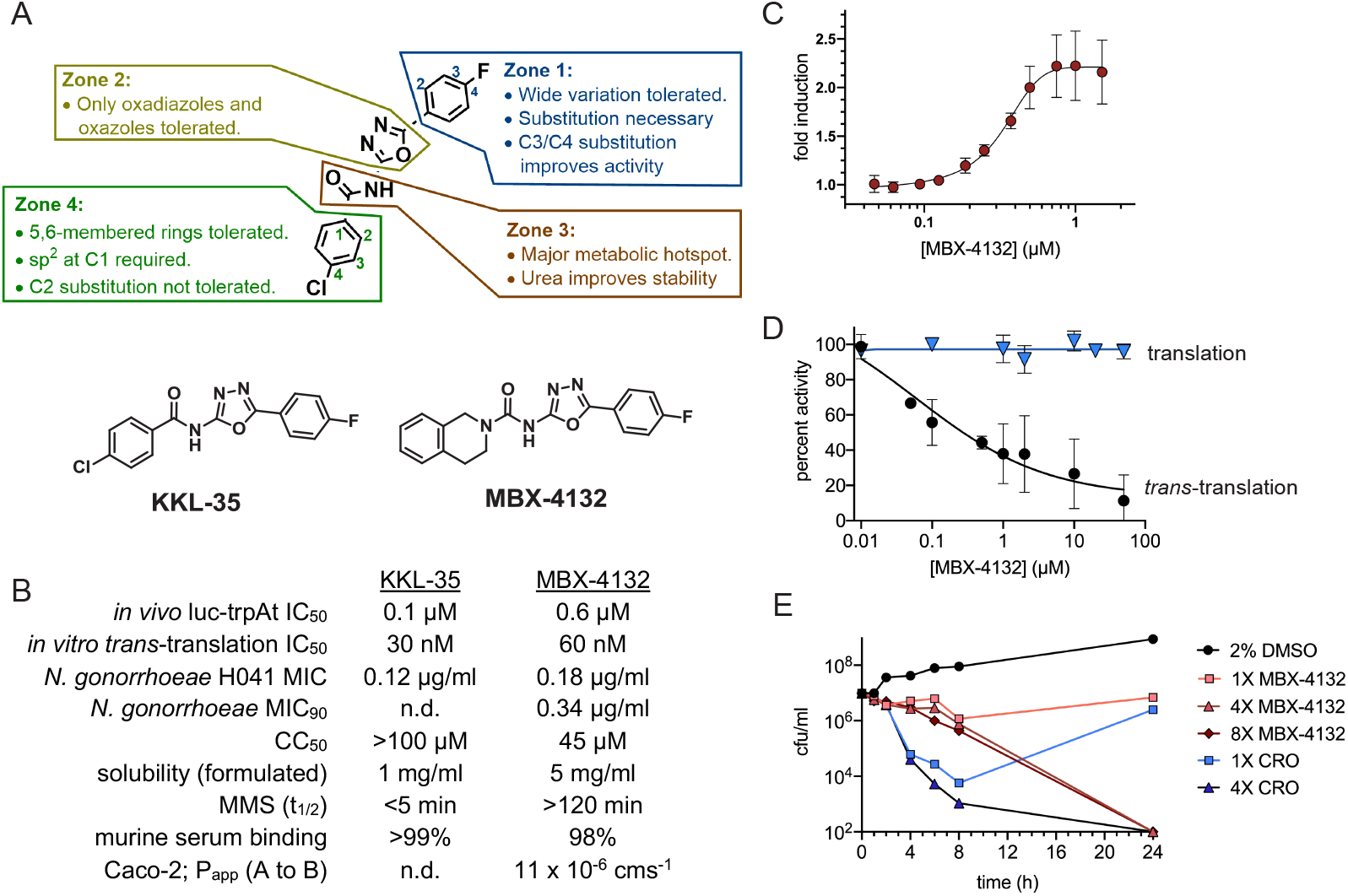
Optimized acylaminooxadiazoles inhibit *trans*-translation to kill *N. gonorrhoeae*. (A) Zones used to guide synthetic strategy with characteristics that govern activity are indicated, and the structure of KKL-35 and MBX-4132 are shown. (B) Properties of the initial hit, KKL-35, and optimized inhibitor MBX-4132 (CC_50_ – half-maximal cytotoxic concentration against HeLa cells; MMS – murine liver microsome stability). (C) Inhibition of *trans*-translation in *E. coli* cells was monitored using a non-stop luciferase reporter. The average of two biological repeats is shown with error bars indicating standard deviations. (D) Inhibition of *trans*-translation in vitro was assayed using an *E. coli* S12 extract to express a truncated, non-stop nano-luciferase gene in the presence of a mutant tmRNA that added the remainder of the nano-luciferase protein. *Trans*-translation activity resulted in luminescence, and addition of MBX-4132 inhibited the reaction (black). As a control, a full-length nano-luciferase gene was used to demonstrate that MBX-4132 does not inhibit translation (blue). The percentage of activity compared to activity in absence of MBX-4132 is shown from the average of at least three repeats with error bars indicating standard deviation. (E) Time-kill assays using *N. gonorrhoeae* show that MBX-4132 is bactericidal at ≥4X MIC. Ceftriaxone (CRO) was used as a control. Counts below the detection limit (100 cfu/ml) were plotted at 100 cfu/ml.

Like the parent acylaminooxadiazole compounds, MBX-4132 exhibited potent broad-spectrum antibiotic activity against Gram-positive species and many Gram-negative species, including *N. gonorrhoeae* (Fig. 1, Tables S3 & S4). The prevalence of multiple-drug resistant (MDR) strains of *N. gonorrhoeae* has made infections increasingly difficult to treat (*2*). MBX-4132 was highly effective against all tested clinical isolates of *N. gonorrhoeae*, including MDR strains (Fig. 1, Table S3), indicating that prevalent resistance mechanisms are not active against MBX-4132. The frequency of spontaneous mutants resistant to MBX-4132 was <1.2 × 10^−9^, suggesting that emergence of resistance in the clinic is likely to be slow. Time-kill assays demonstrated that MBX-4132 was bactericidal to *N. gonorrhoeae* at concentrations ≥ 4X MIC (Fig. 1E).

*In vitro* analyses indicate that MBX-4132 is likely to have low toxicity in mammals. Measurement of competitive binding with 45 mammalian receptors, inhibition of 7 cardiac ion channels, inhibition of the 5 major liver CYP450 enzymes, and an Ames assay for genotoxicity revealed minimal off-target activity, with only minor inhibition of two mammalian receptors observed (Tables S5 & S6). Additionally, high concentrations of MBX-4132 had no effect on mitochondrial membrane polarity in human hepatocytes although elevated levels of reactive oxygen species (ROS) were observed at cytotoxic compound concentrations (Table S7). Likewise, MBX-4132 did not induce differential toxicity against HepG2 cells in the presence of either Glu or Gal, consistent with normal mitochondrial metabolism (Table S7). Collectively, these data show that MBX-4132 is an extremely promising lead candidate suitable for further development.

Pharmacokinetic testing of MBX-4132 revealed that the compound was highly orally bioavailable in mice, exhibiting excellent plasma exposure (area under the curve; AUC), half-life (t_1/2_) and a low clearance rate (Fig. S1). Moreover, animals exhibited no obvious adverse effects at a single dose of 100 mg/kg, or repeat dosing at 10 mg/kg (BID, 7 d). Based on these results and the *in vitro* potency against *N. gonorrhoeae*, we investigated the efficacy of MBX-4132 in a murine genital tract infection model (*12*–*15*). Lower genital tract infection was established with the *N. gonorrhoeae* clinical isolate H041, which is resistant to at least 7 classes of antibiotics (*16*), and mice were treated either with daily intraperitoneal injection of 48 mg/kg gentamicin for 5 days or with a single oral dose of 10 mg/kg MBX-4132 (n = 20–21 mice/group). As previously observed (*13*), daily intraperitoneal injection of gentamicin was effective against H041, clearing infection in 95% of treated mice (Fig. 2A). A single oral dose of MBX-4132 also showed significant efficacy, with 80% of mice completely cleared of infection by 6 days, and a dramatic reduction in bacterial load (Fig. 2). These data are the first *in vivo* proof-of-concept that inhibition of *trans*-translation is a viable antibiotic strategy.

**Figure 2.**
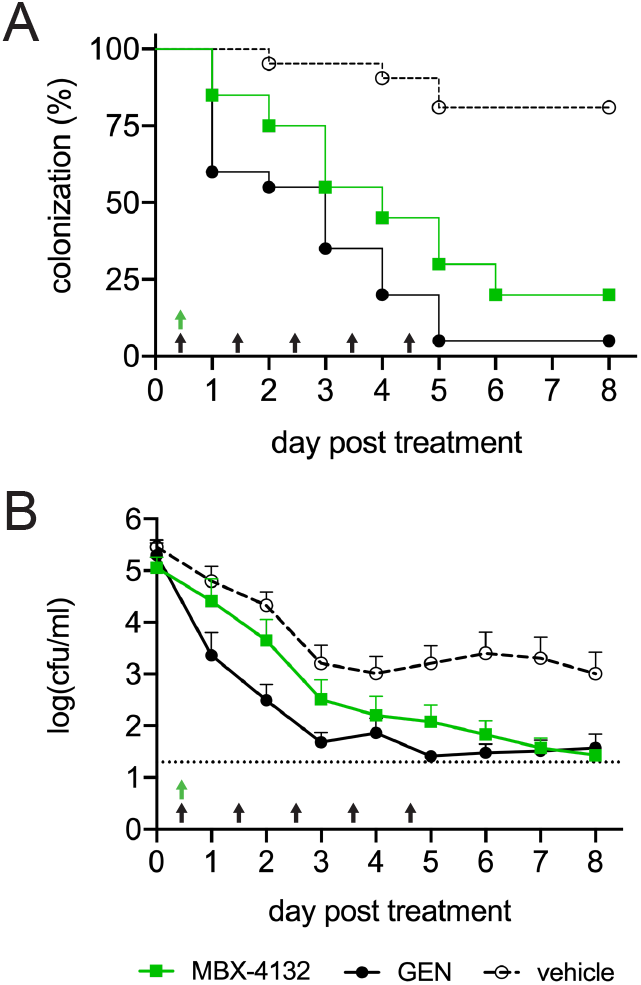
MBX-4132 clears infection by a multiple-antibiotic resistant *N. gonorrhoeae* strain in a murine infection model. Mice were infected with *N. gonorrhoeae* H041 for two days and treated with a single oral dose of 10 mg/kg MBX-4132 or vehicle on day 0 (green arrow), (n = 20 – 21 mice/group). As a positive control, 48 mg/kg gentimicin (GEN) was administered by intraperitoneal injection beginning on day 0 (5QD, black arrows). (A) The percentage of infected mice over 8 days post-treatment. Mice that were culture-negative for at least 3 consecutive days were considered to have cleared infection. MBX-4132 and GEN significantly reduced the percent of infected mice compared to vehicle (*p* < 0.0001). (B) Average bacterial burden (cfu/ml) recovered daily following treatment on day 0. MBX-4132 and GEN significantly reduced the bacterial burden compared to vehicle (*p* < 0.0001). Limit of detection (20 cfu/ml) is denoted by the horizontal dashed line.

To further understand the mechanism of the acylaminooxadiazole antibiotics, we used cryogenic electron microscopy (cryo-EM) to determine the structure of KKL-2098 cross-linked to a non-stop ribosome (Figs. 3 & S2, Table S8). Non-stop ribosomes were generated in *E. coli* by over-expression of an mRNA that contains an RNase III cleavage site before the stop codon and encodes a protein with a histidine tag at the N terminus. Endogenous RNase III cuts this mRNA *in vivo*, and translation produces non-stop ribosomes with the histidine tag from the nascent polypeptide extending from the peptide tunnel. KKL-2098 was added to the culture, UV radiation was used to stimulate cross-linking between KKL-2098 and the ribosomes, and the non-stop ribosomes were affinity-purified, vitrified, and visualized by cryo-EM. 474,382 particles were collected and classified to isolate the 70S ribosomes containing a P-site tRNA, and the structure was solved to 3.1 Å resolution (Figs. S2, S3, S4). Inspection of the map revealed density consistent with KKL-2098 near residue C2452 in the PTC (Fig. S4).

**Figure 3.**
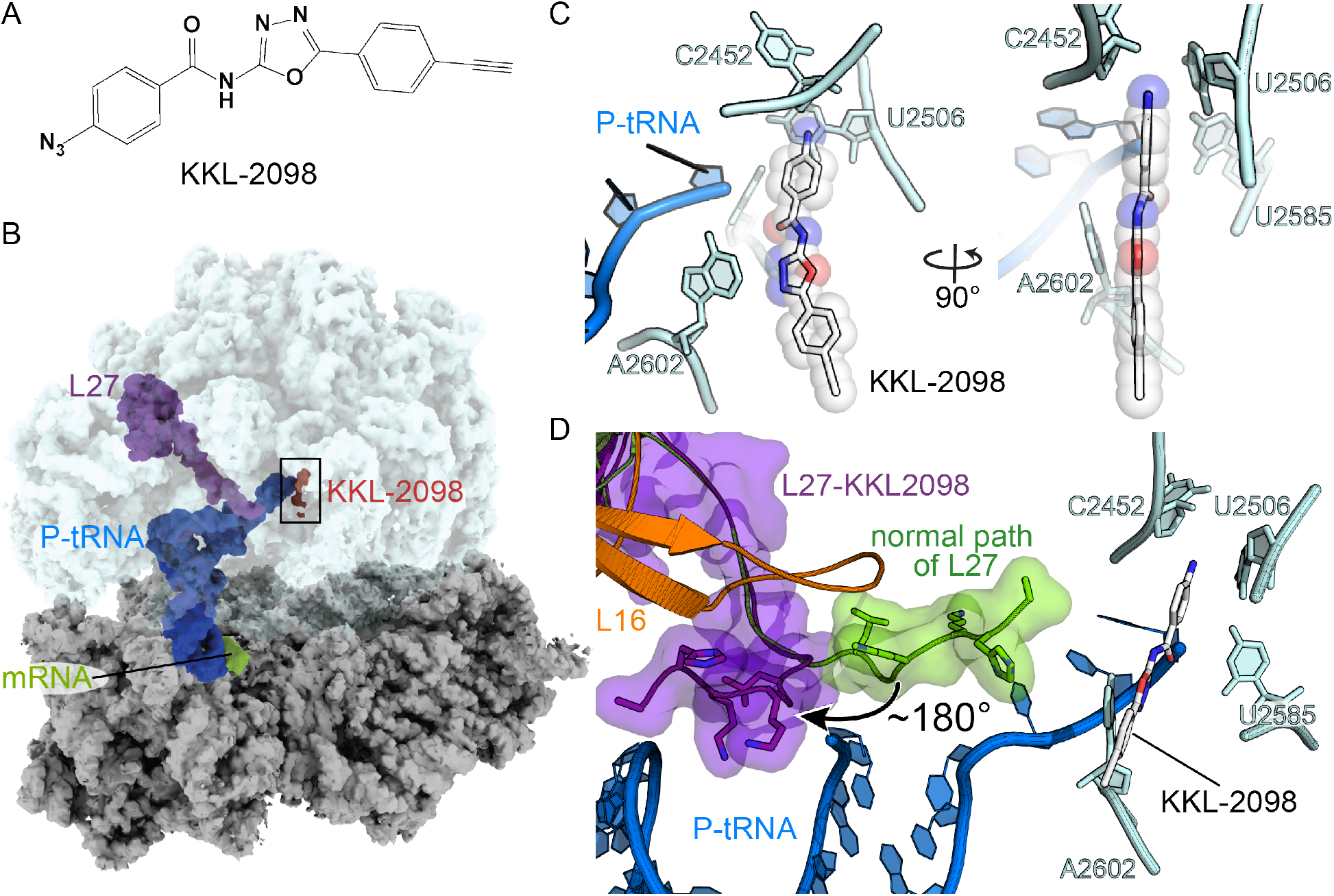
KKL-2098 binds near the peptidyl transferase center. (A) The chemical structure of KKL-2098. (B) The cryo-EM structure of the *E. coli* 70S non-stop ribosome with P-site tRNA, mRNA, ribosomal protein L27 and KKL-2098 indicated. (C) KKL-2098 is positioned close to the peptidyl transferase center adjacent to 23S rRNA A2602, C2452, U2506 and U2585 and the CCA end of the P-site tRNA. (D) The N terminus of L27 (purple) moves 180° to pack against ribosomal protein L16 and the acceptor arm of the P-site tRNA. The normal position of L27 in translating ribosomes is shown in green (PDB ID 6ENU).

KKL-2098 is within 2.1 Å of C2452 (Figs. 3C & S5), the base that was previously shown to crosslink with KKL-2098 in *E. coli* (*3*). The position of KKL-2098 indicates it has hydrophobic interactions with U2506 (Figs. 3C & S5). The amide oxygen is adjacent to the 3’ end of the terminal A76 from the P-site tRNA, and N3 of the oxadiazole is adjacent to A2602, positioned to make hydrogen-bonding or dipole interactions that require the 1,3,4 oxadiazole configuration (note that this oxadiazole configuration has a strong dipole moment (*17*)). These interactions explain why minor changes in Zones 2 and 3 dramatically reduced potency (Table S1). In this position, KKL-2098 overlaps partially with the location of the phosphate of A76 (Fig. S6). The binding site of KKL-2098 is likely to inhibit entry of A-site ligands, including tmRNA-SmpB, into the PTC. Although the binding site of KKL-2098 has some overlap with the adenosine moiety of puromycin, the overall binding sites of KKL-2098 and puromycin are distinct, revealing why these drugs have different mechanisms of action (Fig. S6).

Although no major rearrangements in the rRNA core of the PTC were observed, the N-terminal 7 residues of ribosomal protein L27 move ~180° from the PTC (Fig. 3D). This position is >25 Å from its position in ribosomes containing a peptidyl-tRNA in the P site (*18*–*20*) (Fig. S7). As is the case for many ribosomal proteins, the first 20 residues of L27 form a long extension that lacks secondary structure and thus these residues are frequently not resolved in structures of 70S ribosomes lacking A-site tRNA due to lower resolution. Cross-linking and biochemical studies showed that the N terminus of L27 extends to the 3’ end of the P-site tRNA and stabilizes product formation of the peptidyl transferase reaction (*17*)(*21*–*23*). Consistent with these data, the N terminus has been observed to extend parallel to the 3’ end of the P-site tRNA in structures containing either peptidyl P-site tRNAs or aminoacylated tRNAs at the A site (*18*–*20*). In the KKL-2098-bound structure, the N terminus bends 180° at Gly8 and packs against ribosomal protein L16 and the acceptor stem of the P-site tRNA. These two positions suggest the N terminus of L27 is mobile and its movement may be required for optimal function such as in *trans*-translation or peptide release by any alternative rescue factors. In addition, single-particle reconstruction of non-stop ribosomes lacking a P-site tRNA but containing KKL-2098 also showed the rotated conformation of L27, suggesting that this conformation is preferred when an acylaminooxadiazole is bound (Fig. S7).

To test whether the position of L27 plays a role in acylaminooxadiazole activity, we examined the effects of *E. coli* mutants that are truncated by 3 or 6 residues from the N terminus, preventing L27 from reaching the PTC and the location of acylaminooxadiazole binding. Both mutants were hypersensitive to MBX-4132, but not to other antibiotics that target the ribosome, indicating that absence of L27 from the PTC increases sensitivity to MBX-4132 (Fig. 4).

**Figure 4.**
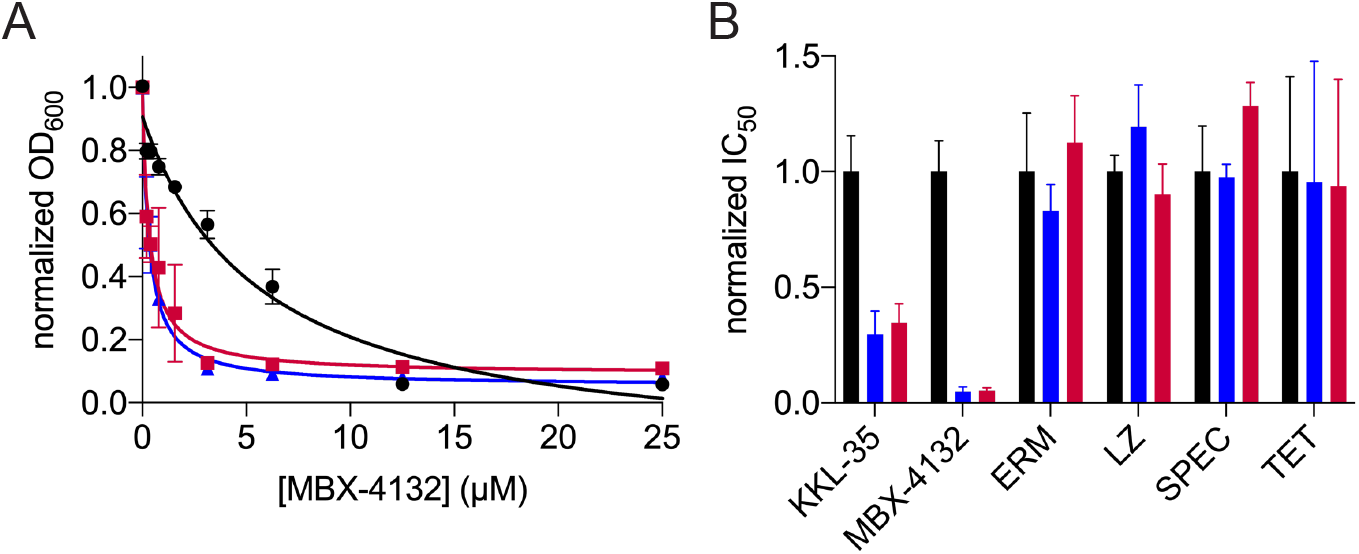
Truncation of L27 causes hypersensitivity to acylaminooxadiazoles. (A) Growth of *E. coli ΔtolC* expressing full-length L27 (black) or variants missing residues 2-5 (−3, blue) or 2-8 (−6, red) was monitored in broth microdilution experiments and the IC_50_ for MBX-4132 was determined. Two technical replicates were performed for each biological replicate and the average of three biological replicates is shown with error bars indicating standard deviations. (B) Normalized mean IC_50_ values from at least 3 biological replicates of experiments as in (A) show that truncation of L27 increases sensitivity to KKL-35 and MBX-4132 but has no effect on erythromycin (ERM), linezolid (LZ), spectinomycin (SPEC) or tetracycline (TET). Error bars indicate standard deviations.

The movement of L27 also provides a possible explanation for why MBX-4132 specifically inhibits *trans*-translation and not translation elongation. Extension of L27 into the PTC in translation elongation complexes would disfavor MBX-4132 binding, but rotation of L27 in non-stop ribosomes could allow MBX-4132 binding and inhibition of *trans*-translation. Although further experiments will be required to test this model, it is clear that the acylaminooxadiazoles exhibit a novel mechanism of antibacterial activity. Taken together, the specific inhibition of *trans*-translation by acylaminooxadiazoles and significant *in vivo* efficacy against *N. gonorrhoeae* after a single oral dose, combined with the new chemical structure, support further development of these compounds as promising new antibiotics.

## ACKNOWLEDGEMENTS

We are grateful to Shura Mankin for providing L27 mutants; Tim Murphy and the staff at Neosome Life Sciences LLC for support in performing and evaluating murine tolerability studies; Charles River Labs, for performing murine pharmacokinetic studies; Micromyx for performing MIC90 studies; Eurofins Panlabs, RTI International and SRI Biosciences for performing extensive *in vitro* safety screening studies. Funding and support for this research was provided by NIH grant R01GM121650 to KCK and by NIAID preclinical contract services, NIH grant R43AI141132 to ZDA, and R01AI132276 to ZDA.

## Notes

### Competing Interest Statement

The authors have declared no competing interest.

