## Supplemental text, methods, tables, figures for "Ribosome rescue inhibitors clear *Neisseria gonorrhoeae in vivo* using a new mechanism"

PDB accession code: 6OM6; EM accession code: EMD-20121

### SUPPLEMENTAL TEXT

#### Structure-Activity Relationships

Compound optimization studies first focused on evaluation of overall structure activity relationships (SAR), resulting in a large data set, a selection of which is shown in Table S1. Analyses focused on the 4 conceptual zones of the molecule (Figure 1A). Although only partial details on SAR data are included here, a more detailed analysis will be the topic of a future publication.

Zone 1 was tolerant of aliphatic, aromatic and heteroaromatic groups (entries 1-3, Table S1; heteroaromatics not shown), but complete replacement of the Zone 1 substituent with a hydrogen atom was not tolerated (entry 4, Table S1). It should be noted that, although incorporating aliphatic groups in Zone 1 moderately increased potency, extensive oxidative metabolism at this site limited further work in this direction (not shown).

The oxadiazole ring (Zone 2) was critical to activity, with only oxadiazoles and oxazoles displaying measurable, if widely varied, activity, despite evaluation of >20 heterocycles (representative examples include entries 1, 5-6, Table S1). This high level of specificity is consistent with hydrogen-bonding and/or dipole-directed interactions with this moiety in the binding site, as these properties vary widely among oxadiazoles (*1*).

Efforts to identify amide isosteres or alternatives in Zone 3 that avoid the observed amidolysis in KKL-35 proved challenging because variation of the amide was poorly tolerated. Of >10 isosteres examined, only analogs containing amides, ureas or *N*-alkyl amides were tolerated (entries 1, 7-9, and 12-16, Table S1). This specificity is consistent with binding interactions involving the oxygen of the Zone 2 amide and a requirement for planarity in this region of the molecule. Although replacement of the amide with a urea prevented amidolysis, simple ureas such as MBX-4346 (entry 12, Table S1) still demonstrated limited microsomal stability, consistent with aliphatic ring oxidation.

Variations in Zone 4 examined both amides and ureas. Among the amides, changes that disrupt coplanarity with the Zone 2 amide were poorly tolerated (*e.g.* aliphatics or C-2 substituents; entries 10 and 11, Table S1). Aromatic groups were well tolerated in Zone 1, with hydrophobic substituents distal to the core leading to improved potency (not shown). Among the ureas, small rings were moderately tolerated, with 6- and 7-membered rings providing improved potency relative to 5 membered rings (entries 12-14, Table S1). Hydrophilic groups, such as the ether moiety in the morpholine ring of MBX-4697 (entry 15, Table S1) had a deleterious effect on potency, although they dramatically increased microsomal stability, consistent with limiting aliphatic group oxidation. Spirocyclic and bridged bicyclic rings had a significantly deleterious effect on potency, consistent with a need for a relatively flat cross-section in this region of the molecule (not shown). Fused bicyclic groups were generally well tolerated, provided they did not include branching on the carbon directly attached to the urea moiety; of these, fused aryl rings provided excellent enhancement in both potency and metabolic stability, resulting in the identification of MBX-4132 (entry 16, Table S1).

Throughout SAR studies, *in vitro* ADME properties such as solubility, microsomal stability and cytotoxicity were monitored. Most compounds exhibited cytotoxicity profiles favorable for development and moderate-to-good solubility, likely due to the hydrophilic 1,3,4-oxadiazole moiety (*1*). An initial concern was metabolic stability, with the Zone 2 amide providing a particular liability. Replacement of this group with a urea moiety provided a clear solution to the problem (entries 12-16, Table S1). Further evaluation of MBX-4132 focused on ligand efficiency, serum effects and permeability (Table S2). These studies revealed excellent drug-like properties and predicted the observed oral bioavailability. It should be noted that MBX-4132 is highly serum-bound, a feature that has an impact on its potency in the presence of serum (Table S2).

### SUPPLEMENTAL METHODS

#### Chemical Synthesis

All commercially obtained reagents and solvents were used as received. $^{1}H$ and $^{13}C$ NMR spectra were recorded on a Bruker 300 MHz instrument. Chemical shifts are given in d values referenced to the internal standard tetramethylsilane (*2*). LC/MS analyses were performed on a Thermo-Finnigan Surveyor LC unit connected to a Thermo LTQ Fleet MS unit. HPLC purification was performed on a Gilson Unipoint instrument equipped with a 00G-4252-P0-AX C18, 10 micron, 150 mm or 250mm x 21.2 mm column from Phenomonex. Silica column purification was performed on Isco brand Combi-flash $R_f$ liquid chromatography system using 50□m silica Luknova SuperSep columns. Melting points were taken on EZ-Melt automated melting point apparatus (Stanford Research Systems, Inc.) in manual mode, and are uncorrected. Thin-layer chromatography was performed on silica gel GHLF plates from Analtech (Newark, DE), and the chromatograms were visualized under UV light at 254 nm. 5-(4-fluorophenyl)-1,3,4-oxadiazol-2-amine, 3-phenyl-1,2,4-oxadiazol-5-amine were purchased from Enamine (Kiev, Ukraine); 5-phenyl-1,2,4-oxadiazol-3-amine was purchased from Chembridge (San Diego, CA, USA); 5-cyclohexyl-1,3,4-oxadiazol-2-amine, 1,3,4-oxadiazol-2-amine were purchased from Life Chemicals (Niagara-on-the-Lake, ON, Canada); p-Toluenesulfonyl Chloride, 1,1'-Carbonyldiimidazole were purchased from Thermo Fisher Scientific (Acros Organics) (New Jersey, USA); 4-Chlorobenzoyl chloride was purchased from Thermo Fisher Scientific (Alfa

Aesar) (New Jersey, USA); 2-amino-5-phenyl-1,3,4-oxadiazole amine, 4-Chlorobenzoic acid, Cyclohexanecarboxylic acid, 4-Chlorobenzaldehyde, Pyrrolidine, Piperidine, Hexamethyleneimine, Morpholine, Methyl Iodide, Sodium Triacetoxyborohydride, Diisopropylethyl amine, Triethylamine, and all solvents were purchased from Millipore-Sigma (St. Louis, MO, USA); 1,2,3,4-tetrahydro-isoquinoline and 4-chlorobenzene sulfonyl chloride were purchased from Combi-Blocks (San Diego, CA, USA); 4-methyl-1,2,3,6-tetrahydropyridine hydrochloride was purchased from Pharmablock (Hatfield, PA, USA); HATU was purchased from GenScript (Piscataway, NJ, USA).

*General Method A:* Heteroaryl amine (1.1eq) was dissolved in NMP (0.29 M) and triethylamine (1.2 eq) stirred for 10 min at room temperature; to this solution, acid chloride (1 eq) was added dropwise via pipette. The reaction mixture was allowed to continue stirring at room temperature for 16-20 h. If the reaction did not reach completion, the mixture was warmed to 50°C for an additional 16-20 h. Upon completion, reaction mixture was added to water, and the resulting precipitate was filtered and dried. Crude precipitate purified with HPLC (5-95% MeCN/$H_2O$ + 0.1% TFA).

*General Method B:* Carboxylic acid (1.1 eq) and HATU/HBTU (1.2 eq) were dissolved in NMP (0.28 M), to which N,N-diisopropylethylamine (1.2 eq) and heteroaryl amine (1 eq) were added. The reaction was stirred at room temperature for 16-20hr. If the reaction did not reach completion, the mixture was warmed to 50°C for an additional 16-20hr. Upon completion, reaction mixture was added to water, and the resulting precipitate was filtered and dried. Crude precipitate purified with HPLC (5-95% MeCN/$H_2O$ + 0.1% TFA).

*General Method C:* To oxadiazole amine (eq) and 1,1'-Carbonyldiimidazole (1 eq) in N-methyl imidazole (0.56 M) was added 3 Å () molecular sieves (1.6mm pellets: ~4 pellets/mmol). Reaction was stirred at room temperature for 3-16 h. Then, the second amine (1.1 eq) was added to the reaction and continued stirring at room temperature for 2-18 h. Upon completion, reaction mixture was added to water, and the resulting precipitate was filtered and dried. Crude precipitate purified with HPLC (5-95% MeCN/$H_2O$ + 0.1% TFA).

KKL-35/MBX-3535: Prepared according to General Method A to provide material consistent with prior reports (*3*). 15.0 mg (7%); mottled tan powder; $^1$H NMR (DMSO): 12.35 (br, 1H), 8.08-8.01 (m, 4H), 7.65 (d, 2H), 7.47 (t, 2H); LC/MS: 318.0 (M+1); mp: >240°C (decomp.); $R_f$: 0.25 (50% EtOAc/hexanes).

MBX-4083: Prepared according to General Method A. 26.0 mg (14%); off-white solid; $^1$H NMR (DMSO): 12.75 (br s, 1H), 8.07-7.99 (m, 4H), 7.67-7.58 (m, 5H); LC/MS: 300.0 (M+1); mp: 182-188°C; $R_f$: 0.64 (50% EtOAc/hexanes).

MBX-3943: Prepared according to General Method A. 12.0 mg (6%); white solid; $^{1}$H NMR (DMSO): 11.77-11.68 (m, 1H), 8.14-7.91 (m, 4H), 7.76-7.53 (m, 5H); LC/MS: 300.2 (M+1); mp: >102°C (slow); $R_f$: 0.65 (50% EtOAc/hexanes).

MBX-3910: Prepared according to General Method A to provide material consistent with prior reports (*4*). 30.0 mg (16%); white solid; $^{1}$H NMR (DMSO): 12.26 (br, 1H), 8.08-7.96 (m, 4H), 7.67-7.62 (m, 5H); LC/MS: 300.2 (M+1); mp: 250-259°C; $R_f$: 0.34 (50% EtOAc/hexanes).

MBX-4370**:** Prepared according to General Method A to provide material consistent with prior reports (*3*). 26.0 mg (28%); White solid; $^{1}$H NMR (DMSO): 11.95 (bs, 1H), 8.02-8.00 (m, 2H), 7.64-7.61 (m, 2H), 2.94 (m, 1H), 2.02-1.98 (m, 2H), 1.77-1.41 (m, 8H); LC/MS: 306.2 (M+1); mp: 204-206°C; $R_f$: 0.72 (5% MeOH/DCM).

MBX-4367: Prepared according to General Method A. 17.0 mg (13%); white solid; $^{1}$H NMR (DMSO): 12.12 (bs, 1H), 9.10 (s, 1H), 8.06-7.98 (m, 2H), 7.65-7.59 (m, 2H); LC/MS: 224.0 (M+1); mp: 212-214°C; $R_f$: 0.32 (5% MeOH/DCM).

MBX-C4227: Material was purchased and tested as received from Life Chemicals, Inc.

MBX-3709: Prepared according to General Method A. 150 mg (97%); light brown solid; $^{1}$H NMR (DMSO): 11.66 (s, 1H), 7.99-7.95 (m, 2H), 7.47-7.41 (m, 2H), 1.87-1.66 (m, 6H), 1.41-1.22 (m, 6H); LC/MS: 290.0 (M+1); mp: >205°C (slow); $R_f$: 0.73 (3.75:46.25:50 MeOH/EtOAc/DCM).

MBX-3776: Prepared in a manner to that previously described (*5*), by dissolving (E/Z)-N-(4-chlorobenzylidene)-5-(4-fluorophenyl)-1,3,4-oxadiazol-2-amine (0.331mmol, 1.0 eq), and $NaHB(OAc)_3$ (0.430 mmol, 1.3 eq) in dichloromethane (1 ml), stirred at room temperature for 18 h. Adsorbed material onto Celite and isolated product from column chromatography eluted with linear gradient of 0-50% EtOAc in Hexanes. 27.0 mg (27%); White solid; $^{1}$H NMR (DMSO): 8.39 (t, 1H), 7.87-7.82 (m, 2H), 7.41-7.34 (m, 6H), 4.43 (d, 2H); LC/MS: 304.1 (M+1); mp: 165-167°C; $R_f$: 0.42 (50% EtOAc/hexanes).

MBX-4076: Prepared by dissolving 4-chloro-N-(5-(4-fluorophenyl)-1,3,4-oxadiazol-2-yl)benzamide (MBX-3535, 0.220 mmol, 1.0 eq) in DMF (2 ml), to which $K_2CO_3$ (0.264 mmol, 1.1 eq) was added and the mixture was stirred at room temperature for 2 h. Then methyl iodide (0.220 mmol, 1.0 eq) was added and continued to stir at room temperature for an additional 65 h. The reaction was diluted with water (~30 ml), solid precipitate was filtered and dried under high vacuum. Product was purified by HPLC (20-100% MeCN/$H_2O$ + 0.1% TFA) and freeze dried to solid. 13.0 mg (18%); White solid; $^{1}$H NMR ($CDCl_3$): 8.21 (d, 2H), 8.01-7.97 (m, 2H), 7.40 (d, 2H), 7.22 (t, 2H), 3.74 (s, 3H); LC/MS: 332.2 (M+1); mp: >148°C (slow); $R_f$: 0.66 (50% EtOAc/hexanes).

MBX-4063**:** Prepared according to modified General Method A; used solvent mixture of Pyridine/Dichloromethane (1/1 mixture) instead of NMP and trietheylamine. 33.0 mg (18%); white solid; $^1$H NMR ($CDCl_3$): 7.96-7.86 (m, 4H), 7.34-7.16 (m, 4H + CHCl3), 2.42 (s, 3H); LC/MS: 334.1 (M+1); mp: 218-221°C; $R_f$: 0.14 (50% EtOAc/hexanes).
MBX-4346**:** Prepared according to General Method C. 15.0 mg (20%); light yellow solid; $^1$H NMR ($CDCl_3$): 7.98-7.95 (m, 2H), 7.20-7.14 (m, 2H), 3.57 (m, 4H), 1.96 (m, 4H); LC/MS: 277.1 (M+1); mp: 197-199°C; $R_f$: 0.94 (10% MeOH/DCM).
MBX-4699: Prepared according to General Method C. 13.0 mg (9%); white solid; $^1$H NMR ($CDCl_3$): 7.97-7.92 (m, 2H), 7.21-7.15 (m, 2H), 3.65 (m, 4H), 1.62 (m, 6H); LC/MS: 291.5 (M+1); mp: 200-203°C; $R_f$: 0.64 (10% MeOH/DCM).
MBX-4700: Prepared according to General Method C. 15.0 mg (33%); white solid; $^1$H NMR ($CDCl_3$ + MeOD): 7.92-7.88 (m, 2H), 7.13-7.07 (m, 2H), 3.48 (m, 4H), 1.69 (m, 4H), 1.52-1.50 (m, 4H); LC/MS: 305.7 (M+1); mp: 191-194°C; $R_f$: 0.63 (10% MeOH/DCM).
MBX-4697: Prepared according to General Method C. 8.9 mg (15%); white solid; $^1$H NMR ($CDCl_3$ + MeOD): 7.93-7.89 (m, 2H), 7.18-7.12 (m, 2H), 3.66 (m, 8H); LC/MS: 293.9 (M+1); mp: 211-216°C; $R_f$: 0.55 (10% MeOH/DCM).
MBX-4366: Prepared according to General Method C. 19.0 mg (23%); white solid; $^1$H NMR ($CDCl_3$): 7.98-7.93 (m, 2H), 7.21-7.16 (t, 2H), 5.41 (m, 1H), 4.11 (m, 2H), 3.78 (m, 2H), 2.11 (m, 2H), 1.73 (s, 3H); LC/MS: 303.0 (M+1); mp: 180-181°C; $R_f$: 0.74 (5% MeOH/DCM).
MBX-4132: Prepared according to General Method C; dried precipitate solids were triturated in MeOH (~50mL) yielded pure product. 375 mg (40%); white solid; $^1$H NMR (DMSO): 7.98-7.94 (m, 2H), 7.46-7.40 (m, 2H), 7.19 (s, 4H), 4.69 (s, 2H), 3.75 (t, 2H), 2.85 (t, 2H); LC/MS: 339.1 (M+1); mp: >190°C (slow); $R_f$: 0.51 (50% EtOAc/hexanes).

**Bacterial Strains and Growth Conditions**

Bacterial strains, plasmids, and synthetic sequences are shown in Table S9. *E. coli* strains expressing L27 were constructed by transducing a *tolC::cat* allele into IW312 strains using a P1 *vir* phage. These strains were grown in LB medium containing 1 mM IPTG. To quantify the *in vitro* antibacterial activity of acylaminooxadiazole analogs against various bacterial strains, the minimum inhibitory concentration (MIC) was measured using microbroth dilution assays as described in the CLSI guidelines (M7-A7) (*6*), except that liquid G77L medium (*7*) was used for *Neisseria gonorrhoeae* MIC assays. Each assay comprised three technical replicates, and each assay was repeated at least once. The reported MIC is the geometric mean of at least 5 technical replicates. H041(STM$^R$) is a streptomycin-resistant derivative of the multi-drug resistant (MDR), ceftriaxone-resistant strain H041 (Ohnishi) and was cultured as previously described (*8–10*).

**Time-kill assays**

The time kill assay was performed essentially as described (*11*) with the following modifications for *N. gonorrhoeae*. The bacterial inoculum for the assays was prepared by suspending colonies of *N. gonorrhoeae* ATCC 49226 grown on a chocolate agar plate for >24 h (37 °C with 5% $CO_2$) in G77L medium. The cell suspension was adjusted to an $OD_{600}$ of 0.1 and was diluted 1:10 in G77L media (final cell density ~1 × $10^7$ cells/ml) containing various concentrations of MBX-4132. The resulting cultures were incubated at 37 °C (5% $CO_2$), and viability was monitored over 24 h by removing samples at various time points, making serial 10-fold dilutions in G77L, and spotting 5 µl of each dilution onto the surface of a chocolate agar plate in triplicate. Colonies were counted after the plates were incubated at 37 °C (5% $CO_2$) for 18-18 h, colony forming units (cfu) per ml were calculated, and the average and standard deviation for the three replicates was determined. The lower limit of detection of this assay was determined to be 100-200 cfu/ml. This experiment was repeated three times, and the results of a representative experiment are shown.

***In vitro trans*-translation and translation assays**

The *in vitro trans*-translation assay is designed to produce the first 157 amino acids of nano-luciferase from a non-stop mRNA (nanoluc-ns) and add the C-terminal peptide AAVSGWRLFKKIS via a mutant version of *E. coli* tmRNA (tmRNA-nl) to reconstitute an active nano-luciferase (*12*). In this assay, the tagged nano-luciferase gene has >30,000-fold increase in activity over the peptide made in the absence of tmRNA-nl.

S12 lysates were made according to the procedure of Kim, et al., (*13*). Briefly, a culture of *E. coli* BL21 (DE3) cells containing a pET28 plasmid carrying the *E. coli* T7 polymerase gene was grown at 37 °C to an $OD_{600}$ = 0.8, induced with 1 mM IPTG and grown for an additional 3 h. Cells were harvested by centrifugation at 20,000 *g* for 10 min at 4°C and the pellet was resuspended in buffer A (20 mM Tris-acetate (pH 8.2), 14 mM $Mg(OAc)_2$, 60 mM potassium glutamate, 1 mM DTT). Cells were lysed by sonication, the lysate was clarified by centrifugation at 12,000 *g* for 10 min, and the supernatant was stored at -80 °C.

*E. coli* SmpB was purified as previously described (*14*). tmRNA-nl, a variant of *E. coli* tmRNA encoding the final 11 residues of nano-luciferase, was transcribed from the tmRNA-nl synthetic DNA sequence in vitro, purified, and folded as previously described for wild-type *E. coli* tmRNA (*14*). The nanoluc-stop template DNA was prepared as previously described (*14*) by PCR amplification from pMC1 using T7 universal and nanoluc-stop primers. The nanoluc-ns was prepared in a similar manner using pMC1 with T7 universal and nanoluc-ns primers.

For *in vitro trans*-translation, tmRNA-nl and SmpB were premixed and stored on ice. S12 lysate (2 μl), freshly made polymix buffer (2 μl) (*15*) (final reaction concentrations 5 mM Hepes pH 7.6, 5 mM $NH_4Cl$, 0.5 mM $CaCl_2$, 1.5 mm $MgCl_2$, 1 mM DTT, 8 mm putrescene, 2 mm ATP, 2 mM GTP, 1 mM CTP, 1 mM UTP, 0.3 mM each amino acid, 3 mg/ml *E. coli* tRNAs), nanoluc-ns template (0.5 μl; 60 ng) and 2 μl water were mixed and incubated at 37 °C for 5 min. This solution was dispensed to tubes containing 0.5 μl of different concentrations of an inhibitor prepared in 75% acetonitrile, 25% water, and incubated at 37 °C for 5 min. The tmRNA-nl/SmpB mixture (0.5 μl each, 2 μM final) was added to each tube and the samples were incubated at 37 °C for 1.5 h. NanoLuc substrate (Promega) was prepared according to the manufacturer's instructions, one volume of this substrate solution was added to each sample tube, and the reactions transferred to a white 96-well plate. Luminescence readings were obtained using the SpectraMax i3 microplate reader. Data analysis was performed using GraphPad Prism 8.

**Frequency of resistance**

First, MIC values were determined for MBX-4132 using the reference agar dilution method (*6*, *16*). *N. gonorrhoeae* strain 49226 (ATCC) was suspended to the equivalent of a 5 McFarland standard in G77L broth and then diluted to generate the final inoculum ($1.5 \times 10^5$). The bacterial cell suspension was then transferred to wells in a stainless-steel replicator block which was used to inoculate the test plates. After the inoculum had dried, all plates were incubated at 35°C in 5% $CO_2$. The MIC was read post-incubation per CLSI guidelines (*6*, *16*). To determine the frequency of resistance, stock solutions of MBX-4132 were prepared at 100X the final test concentrations of 4× and 8× the predetermined MIC value. A 0.5 mL aliquot of the 100X stock was mixed with 49.5 mL of molten GC Medium agar/1% IsoVitaleX to produce an agar/drug mixture that was either 4- or 8-fold the MIC and dispensed into sterile 150 x 15 mm plates (VWR) at a volume of 50 mL per plate. A dense cell suspension equivalent to 5 McFarland was prepared using bacterial growth from 48 h chocolate agar plates of *N. gonorrhoeae* (ATCC 49226). The viable count of each suspension was determined by plating serial ten-fold dilutions onto GC Medium agar/1% IsoVitaleX in duplicate. A 0.25 ml aliquot of inoculum was spread onto the surface of duplicate 150 x 15 mm test plates. After allowing the inoculum to dry on the surface of the plate, the plates were inverted and incubated at 35ºC (with 5% $CO_2$) for 48 h. Colony counts were determined manually and the spontaneous mutation frequency was calculated using the following equation: Average number of colonies from selection plates/Total number of cells inoculated

If there were no colonies on the antibiotic selection plates, the spontaneous mutation frequency was calculated as 1/inoculum to indicate that the spontaneous mutation frequency was less than the limit of detection (one cfu).

**Mammalian cell cytotoxicity ($CC_{50}$)**

The half maximal cytotoxic concentration ($CC_{50}$) of each compound against HeLa cells (ATCC CCL-2) was measured as previously described (*17*). Each assay comprised three technical replicates, and each assay was repeated at least once. The mean values for each biological replicate were averaged, and the $CC_{50}$ was determined using a 4-parameter nonlinear curve fitting algorithm (GraphPad Prism). The average of the $CC_{50}$s from two biological replicates was calculated and reported.

**Murine liver microsome stability**

To examine potential for first-pass metabolism of analogs in the liver, the stability of analogs in the presence of mouse liver microsome preparations (Eurofins Discovery for human, dog and rat; Xenotech for mouse) was measured using the method of Kuhnz, et al. (*18*) for murine studies and Oback for the dog, human and rat studies. (*19*). The amount of parent compound remaining after incubation with microsomes in the presence of NADPH over a 30 min time range was measured using a reverse-phase liquid chromatography/mass spectroscopy method that was customized for each compound. Half-lives were calculated using linear regression analysis of several time points.

**Caco-2 permeability**

To evaluate the potential for oral bioavailability, the ability of prioritized compounds to permeate a monolayer of Caco-2 intestinal epithelial cells was determined as described (*20*). Caco-2 permeability values ($P_{app}$) >1 × $10^{-6}$ cm/sec are predictive of oral bioavailability. The observation that $P_{app\ A\rightarrow B} > P_{app\ B\rightarrow A}$ indicates that efflux from the basolateral compartment does not occur.

**Serum Protein Binding**

Serum protein (fetal bovine) binding was determined using an equilibrium dialysis method as described (*21*). The amount of compound in each chamber (buffer and serum) was measured using methods for reverse-phase liquid chromatography/mass spectroscopy methods that were customized for each compound.

**Aqueous solubility**

The maximum aqueous solubility of each compound was determined using a nephelometric method as described (*22*). Each assay comprised three technical replicates, and each assay was repeated at least once. The reported solubility is the average of at least 5 technical replicates.

**Cell-based non-stop luciferase reporter assay**

To verify that MBX-4132 retains activity as an inhibitor of trans-translation, we measured its dose-dependent activity against the non-stop luciferase reporter assay strain E. coli SB75 ΔtolC::kan (pluc-trpAt) essentially as described (*14*), with modifications. Briefly, serial 1.5-fold dilutions of MBX-4132 in DMSO were transferred to 96-well assay plates (Costar 3195), followed by the addition of 50 µl of an overnight culture of SB75 ΔtolC::kan (pluc-trpAt) that had been diluted to a final $OD_{600}$ of 0.4 with LB media supplemented with 100 µg ampicillin/ml and 1 mM IPTG. The final concentrations of MBX-4132 ranged from 0.05 – 1.5 µM, and the final concentration of DMSO was 2%. The assay plates were incubated at room temperature for 2 h, and 50 µl of BrightGlo (Promega) bioluminescence reagent was added to each well. After 10 min incubation, bioluminescence intensity was measured using an Envision multi-label plate reader (Perkin Elmer). Each assay comprised three technical replicates, and the experiment was repeated three times. The fold induction for each MBX-4132 treated sample as compared to the DMSO-only sample was calculated for each technical replicate, and the average and standard deviation of the three technical was calculated. The $IC_{50}$ was determined using a 4-parameter nonlinear curve fitting algorithm (GraphPad Prism).

**CYP450 Inhibition, Receptor Panel Profiling, and Cardiac Ion Channel Profiling**

Several *in vitro* selectivity assays were performed at Eurofins Discovery Services using established methods and controls that behaved as expected (Tables S5 & S6). For CYP450 inhibition assays, activity of MBX-4132 was tested at 5 concentrations from 30 nM to 100 µM; no inhibitory activity >50% was observed at any concentration. For receptor panel profiling, MBX-4132 was evaluated at 10 µM and activity of >50% (agonist or antagonist) was scored as active.

**Ames Assay**

The Ames assay was performed at SRI Biosciences, following the standard protocols established there (*23–25*). In brief, samples were evaluated for their ability to induce genetic damage using the plate incorporation method with *Salmonella typhimurium* strains TA98 and TA100 with and without a metabolic activation mixture containing 10% Aroclar-1254-induced rat-liver microsomes

(S9). MBX-4132 was tested from a 5 mg/mL DMSO stock solution (the solubility limit), which provided plate concentrations up to 500 μg/plate, with serial dilutions accessing doses as low as 5 μg/plate. MBX-4132 precipitated at the two highest dose levels (500 and 100 μg/plate), but at 50 μg/plate, no precipitation was observed. Precipitation did no interfere with colony counts or analysis. Test articles were considered mutagenic when the mean number of revertant colonies increased in a dose-dependent manner. Some cytotoxicity was observed, but sufficient colonies were formed to allow analysis. As shown in Table S6, MBX-4132 exhibited no significant deviation in revertant colonies from the DMSO control at any concentration tested. Data for 2-nitrofluorene and sodium azide (positive controls) and DMSO (negative control) were included for reference.

**Mitochondrial Toxicity Assays (performed at Eurofins Cerep Panlabs)**

*Multiplexed cytotoxicity assay*

Human primary hepatocytes were grown on collagen I coated optical plates, cultured in hepatocytes culture media in a humidified 5% $CO_2$ atmosphere at 37 ˚C. Cells were incubated in the presence of MBX-4132 at 10 concentrations starting a 100 μM and serially diluted 3.16-fold for 24 h at 37 °C, incubated for 30 min with multiplexed fluorescent dyes (Hoescht, 6-carboxy-2',7'-dichlorodihydrofluorescein diazetat, and TMRE) and imaged to allow visualization of nuclei, reactive oxygen species generation, and mitochondrial oxidation. A >3.5-fold induction of ROS was considered consistent with formation of ROS and a >2-fold change in TMRE signal indicated an increase or decrease in mitochondrial membrane potential (*26*).

*Mitochondrial toxicity assay*

HepG2 cells were seeded and cultured in media containing either glucose or galactose overnight. Test compounds were added at 8 concentrations (3-fold serial dilution from 100 μM to 30 nM) and incubated with the cells for 24 h. Cell viability was measured by the alamarBlue method. The data in Table S7 are inconsistent with MBX-4132 disrupting mitochondrial metabolic processes (*27*).

**Pharmacokinetic analyses in mice**

*For PO suspension dosing* (Performed at Neosome LLC):

Female CD-1 mice were fasted for 2 h prior, and 4 h after dosing. MBX-4132 was administered male BalB/C mice at 10 mL/kg via oral gavage with a 1.0, 2.5 or 10.0 mg/ml suspension in vehicle A (5% DMSO, 5% Cremophor EL®, 0.45% hydroxypropylmethylcellulose, 0.45% alginic acid, 22.5% hydroxy-betacyclodextrin). At 0.25, 0.5, 1, 2, 4, 8 and 24 h post dose, 3 mice from each group were euthanized by $CO_2$ inhalation, blood was collected by cardiac puncture into $K_2$EDTA collection tubes, and protein was precipitated and analysed by LC/MS-MS for plasma concentrations of MBX-4132. Data were analysed using WinNonLin.

*For IV and SC dosing* (performed at Charles River Labs)*:*

MBX-4132 was administered at 6 ml/kg via direct tail vein puncture (IV, slow push), oral gavage (PO) or subcutaneously to the intrascapular region (SC) in a 2 mg/ml 10% DMSO/80% PEG-400/10% water formulation. For oral dosing, mice were fasted for 2 h prior and four hours post dosing. At 0.083, 0.5, 1, 4, 8 and 24 h post dose, blood was collected from 3 mice/group by tail vein or facial bleed into $K_2EDTA$ collection tubes, and protein was precipitated and analysed by LC/MS-MS for plasma concentrations of MBX-4132. Sampling was performed serially, with each mouse contributing sample at all time points. Data were analysed using WinNonLin.

**Murine tolerability studies**

(All testing was performed at Neosome LLC)

*Single dose tolerability studies:*

Female CD-1 mice were fasted for 2 h prior, and 4 h after dosing. MBX-4132 was administered at 10 ml/kg via oral gavage with a 0.0 (vehicle control), 1.0, 2.5 or 10.0 mg/ml suspension in vehicle A. Mice were observed at 0.083, 0.25, 0.5, 1, 2, 5, 8 and 24 h post dosing, and any abnormal observations (behaviour, agility, coat condition and appearance, color of urine, quality of feces, etc.) were noted. No abnormal observations of any dosing group were made during the 24 h course of this study.

*Multidose tolerability studies:*

In preparation for this study, an abbreviated version of the above murine PK study using the same formulation examined fasted mice vs. fed mice was performed, with plasma samples taken at 1 and 4 h post dosing and analyses as described in the PK section. No significant variation in exposure was observed at either timepoint.

Female CD-1 mice were given free access to food and water. MBX-4132 was administered to groups of 3 mice for 7 d either QD (with compound) or BID (two groups, one vehicle only and one with compounds) at 10 ml/kg via oral gavage with a 1.0 mg/ml suspension in A. Mice were observed twice daily for 10 d (3 d post final dose), additionally, mice were weighed daily. Any abnormal observations (behaviour, agility, weight, coat condition and appearance, color of urine, quality of feces, etc.) were noted. No abnormal observations of any dosing group were made during the 24 h course of this study. One mouse in the vehicle only group did exhibit a slight weight loss but had recovered weight by the end of the study.

#### *In vivo* efficacy testing in the gonorrhea mouse model

Groups of female BALB/cAnNCr mice (Charles River Laboratories) (6-7 weeks old) were treated with 17β-estradiol and antibiotics (streptomycin and trimethoprim) to increase susceptibility to long-term *N. gonorrhoeae* infection as described (*8*). Mice were inoculated vaginally with *N. gonorrhoeae* strain H041 ($10^4$ cfu/mouse) two days after estradiol pellet implantation and vaginal swabs were cultured for two days post-bacterial inoculation to confirm infection. On the afternoon of the second culture day (day 0), mice were given MBX-4132, GEN or the vehicle (n= 20-21 mice/group). Doses of MBX-4132 were prepared fresh in vehicle at the time of treatment and administered as a single oral dose (dose volume 10 ml/kg). The positive control GEN (48 mg/kg) was prepared and administered intraperitoneally as 5 daily doses as previously described (*8*). Vaginal swabs were collected on 8 consecutive days following treatment and quantitatively cultured for *N. gonorrhoeae* to assess efficacy. The data are expressed as CFU/ml of vaginal swab suspension. Clearance was shown by Kaplan-Meier curves with log-rank (Mantel-Cox) statistical analysis. The average cfu/ml over time was compared by 2-way ANOVA with Bonferroni post-hoc analysis. Statistics were performed in GraphPad Prism Software. At the study endpoint (10 days post-inoculation), mice were euthanized using compressed $CO_2$ gas in a $CO_2$ gas chamber in the Laboratory Animal Medicine Facility. All animal experiments were conducted at the Uniformed Services University of the Health Sciences, a facility fully accredited by the Association for the Assessment and Accreditation of Laboratory Animal Care, under a protocol that was approved by the university's Institutional Animal Care and Use Committee.

#### Purification of stalled ribosome complexes

To construct pET28-H10arfArnc, a DNA cassette encoding 10 histidine residues, the M2 epitope, 3 glycine residues, and 20 arbitrary codons followed by 71 base pairs from the 3’ end of the *E. coli arfA* gene was synthesized and assembled into pET28 that had been digested with NcoI and HindIII (*28*). The sequence from *arfA* contains the RNase III cleavage site (*29*, *30*). Translation of the cleaved mRNA produces a 59 amino acid peptide with 10 histidines at the N terminus.

*E. coli* 70S ribosomes were purified as described previously (*31*). *E. coli* BL21(DE3) pET28-H10arfArnc cells were grown to an $A_{600}$ of ~0.5 in Luria broth (LB) medium at 37 °C and induced with 1 μM IPTG and 1 μM KKL-2098 (synthesized as previously described (*32*)) and continued to grow for an additional hour at 37 °C then cooled on ice for 20 min. All centrifugation steps were performed at 4 °C. Cells were pelleted by centrifugation and washed with buffer 1 (10 mM HEPES-KOH, pH 7.6, 10 mM $MgCl_2$, 1 M $NH_4Cl$, 6 mM β-mercaptoethanol (β-Me)) twice and then resuspended in buffer 2 (10 mM HEPES-KOH, pH 7.6, 10 mM $MgCl_2$, 100 mM $NH_4Cl$, 6 mM β-

Me). The cells were crosslinked under ultraviolet light (254 nm) for 10 min, then lysed using an EmulsiFlex-C5 high-pressure homogenizer (Avestin). Cell debris was removed by centrifuging at 13,000 × *g* for 15 min. The lysate was further centrifuged at 27,000 × *g* for 30 min to obtain the S30 fraction. Ribosomes were pelleted by centrifuging at 42,000 × *g* for 17 h. The pellets were resuspended in buffer 2 and bound to a 1 mL IMAC gravity column, washed with 10 column volumes of Buffer 2 and eluted with Buffer 2 supplemented with 500 μM imidazole. Ribosomes were further purified over a 10–40% sucrose gradient in buffer 2 at 70,000 × *g* for 12 h. 70S ribosomes were separated from polysomes and subunits using a Brandel gradient fractionator. The 70S fractions were pooled, pelleted, resuspended in buffer 2, and stored at −80 °C.

**Cryo-Electron Microscopy**

UltrAuFoil® grids (Quantifoil, R1.2/1.3) were glow-discharged for 20 s with a Solarus 950 (Gatan). 3 μl of 70S complexes at 100 nM were placed on grids at 8 °C in 100% humidity and blotted for 3.5 s using a Vitrobot Mark IV (FEI).

Two independent datasets of 2,394 micrographs total (1863 and 531 micrographs, respectively) were collected on a Titan Krios (FEI) microscope operated at 300 kV with a C2 aperture diameter of 70 μm. Movie frames were recorded at an accumulated dose of 58 $e^{-}/Å^{2}$ at a magnification of 59,000X (corresponding to a pixel value of 1.191 Å) with a DE-64 direct electron detector in counting mode (*33*) using Leginon (*34*) for automatic data acquisition. Images were recorded with a total exposure time of 19.3 s, and intermediate frames were recorded every 0.2 s giving a total of 78 frames per image. Two independent datasets of 70S ribosomes specimens were collected with an overall of 2,394 micrographs (1863 and 531 micrographs from the first and the second datasets, respectively). Defocus values ranged from -1.3 to -3 μm. All the pre-processing steps were performed in the Appion (*35*).

**Image processing**

**Pre-processing**

All processing steps were carried out using Appion (*35*). All frames of each micrograph were aligned using MotionCor2 (*36*). Contrast transfer function (CTF) parameters were estimated on all motion-corrected micrographs using CTFFIND4 (*37*) and GCTF (*36*) and the best estimate chosen using resolution evaluation in Appion (*38*). 197 micrographs were excluded after manual visualization of their corresponding power spectra displayed ice contamination, exposure to the shifted beam, or low-resolution Thon ring profiles. An initial set of ~2000 particles were picked using DoG (Difference of Gaussian) Picker (*39*). A rotational average was generated from these picks, and this was used as a template for template-based picking using FindEM (*40*). A total of 474,382 particles (373,845 and 100,537 particles from the first and the second dataset, respectively) were picked. Particles were extracted with a box size of 384x384 pixels in Appion.

Extracted particles from the two datasets were processed independently and combined after the last round of 3D reconstruction, and subjected to further 3D classifications and refinements in RELION-3 (*41*) (Figure S2). Processing steps were initially performed on a 4X-binned dataset.

Initial 3D refinement occurred against a 60 Å low-pass filtered empty *E. coli* 70S ribosome from a previous dataset. 3D classification without alignment was used to discard free 50S subunits. The resulting 70S classes were combined and refined using an initial angular sampling of 7.5° and local angular sampling of 1.8° to improve angular assignment. Following refinement, another round of 3D classification without alignment was performed, with low resolution particles and particles containing E-site tRNA discarded. Two classes with an unrotated 70S were combined and subjected to refinement, followed by focused classification with a P-site mask. The P-site mask was generated from a 70S ribosome with P-site tRNA (PDB ID 4V4I) (*42*), with a 5-voxel expansion and a 7-voxel soft edge. This resulted in a class of 70S particles with P-site tRNA. These particles were unbinned, refined, then underwent focused classification with an A site mask, created as stated, using a model of the 70S ribosome with A-site tRNA (PDB ID 4V5D) (*43*). Classes with P-site tRNA but no A-site tRNA were refined and post-processed.

**Post-processing and beam-tilt correction**

The resultant map was post-processed in RELION using a solvent mask generated from the final reconstruction low-pass filtered to 40 Å with a 7-pixel extension and 10-pixel soft edge. Beam-tilt estimation and correction was performed (without per-particle refinement of CTF parameters) using CTF refinement followed by 3D reconstruction. These steps were repeated iteratively until the highest resolution was achieved, and the final map post-processed with a solvent mask. Resolution estimations were calculated from Fourier shell correlations (FSC) at 0.143 between the two independently refined half-maps. Maps were sharpened in PHENIX (*44*). The graphs of directional 3D FSC and global resolution of the maps were plotted using 3DFSC Processing Server (*45*). Local resolution was estimated using blocres in Bsoft (*46*).

### SUPPLEMENTAL TABLES

Table S1: Properties of acylaminooxadiazoles (subset of SAR analogs).

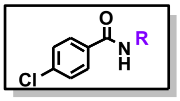

| # | CMPD# | R | *Luc* $IC_{50}$ a | Δ*tolC* *E. coli* MIC[b] | *Ng* MIC[b] | $CC_{50}$ [c] | Sol[d] | MLMS[e] |
|---|---|---|---|---|---|---|---|---|
| 1 | KKL-35/ MBX-3535 | 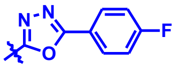 | 0.23 | 0.49 | 0.12 | >100 | 25 | <5 |
| 2 | MBX-3910 | 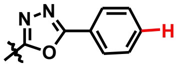 | 0.14 | 1.9 | 0.30 | 40 | 13 | 5 |
| 3 | MBX-4370 | 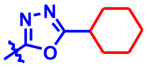 | 0.13 | 0.48 | 0.12 | 17 | 50 | <5 |
| 4 | MBX-4367 | 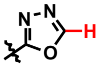 | >25 | 22.3 | 11.2 | >100 | >200 | 8 |
| 5 | MBX-4083 | 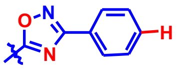 | 1.4 | 2.7 | 4.8 | >100 | 6.3 | <5 |
| 6 | MBX-3943 | 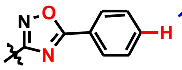 | >25 | >30 | 15.0 | >100 | 13 | <5 |

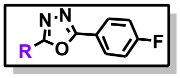

| # | CMPD# | R | *Luc* $IC_{50}$ [a] | Δ*tolC* *E. coli* MIC[b] | *Ng* MIC[b] | $CC_{50}$ [c] | Sol[d] | MLMS[e] |
|---|---|---|---|---|---|---|---|---|
| 7 | MBX-3776 | 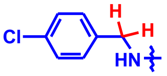 | >25 | >30 | >30 | >100 | 25 | 5 |
| 8 | MBX-4076 | 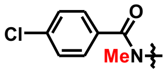 | 1.05 | 4.6 | 2.7 | 95 | <3.1 | |
| 9 | MBX-4063 | 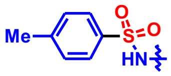 | >25 | >33 | >33 | >100 | 100 | 53 |
| 10 | MBX-3709 | 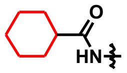 | 12.5 | 28.9 | 4.6 | >100 | 25 | <5 |
| 11 | MBX-C4227 | 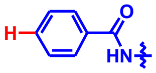 | 1.0 | 0.88 | 1.1 | >100 | 50 | |
| **Urea Variants** | | | | | | | | |
| 12 | MBX-4346 | 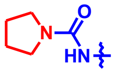 | 14.3 | 27.6 | 4.4 | >100 | >200 | 25 |
| 13 | MBX-4699 | 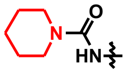 | 2.1 | 7.3 | 0.58 | >100 | 50 | 12 |
| 14 | MBX-4700 | 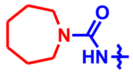 | 1.8 | 4.8 | 0.61 | >100 | 25 | 10 |
| 15 | MBX-4697 | 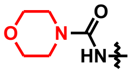 | 9.2 | 29.2 | 4.7 | >100 | 100 | >120 |
| 16 | MBX-4132 | 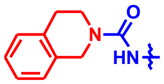 | 0.19 | 2.7 | 0.18 | 45 | 100 | >120 |

[a] Half Maximal activity in Δ*tolC* *E. coli* luciferase assay (μM) [b] MIC vs *Neisseria gonorrhoeae* (49226) or E. coli KLE701 in μg/mL. [c] Against HeLa cells (μM). [d] Solubility in water (μM; nephlometry). [e] Murine liver microsome stability, $t_{1/2}$ in min at 37 °C in the presence of NADPH.

Table S2: In Vitro ADME Properties

| | Microsomal Stability ($t_{1/2}$; min) | | | | serum | | Serum Binding (% bound) | | | | Caco-2 ($P_{app}$; x $10^{-6}$ $cms^{-1}$) | |
|---|---|---|---|---|---|---|---|---|---|---|---|---|
| | Murine | Rat | Dog | Hum. | Shift[a] | Stab[b] | Murine | Rat | Dog | Human | A→B | B→A |
| MBX-4132 | >120 | >120 | >120 | >120 | 12 | >99.8 | 98.0 | 93.0 | 99.0 | 99.0 | 11.1 | 7.3 |

[a]Ratio of MIC +/- 10% fetal bovine serum added. [b]% remaining after 1 h incubation at 37 ˚C.

Table S3: Anti-gonococcal spectrum

| Strain | Resistance/Description | MIC (µg/ml) KKL-35 | MIC (µg/ml) MBX-4132 |
|---|---|---|---|
| ATCC 49226 | type strain | 0.12 | 0.13 |
| ATCC 700719 | SPT | 0.25 | 0.21 |
| ATCC 700825 | STR | 0.06 | 0.04 |
| BAA-1846 | TET | 0.06 | 0.04 |
| HO41 | PEN, TET, CFM, CRO, LVX | 0.25 | 0.17 |
| CDC-0165 | CIP, PEN, TET | n.d. | 0.68 |
| CDC-0166 | CIP, PEN, TET | n.d. | 0.17 |
| CDC-0167 | | n.d. | 0.13 |
| CDC-0169 | CIP, PEN, TET | n.d. | 0.17 |
| CDC-0170 | CIP, PEN, TET | n.d. | 0.17 |
| CDC-0171 | CIP, PEN, TET | n.d. | 0.17 |
| CDC-0172 | CIP, PEN, TET | n.d. | 0.17 |
| CDC-0173 | CIP, PEN, TET | n.d. | 0.17 |
| CDC-0174 | CIP, PEN, TET | n.d. | 0.17 |
| CDC-0175 | CIP, PEN, TET | n.d. | 0.08 |
| CDC-0176 | | n.d. | 0.34 |
| CDC-0177 | CIP, PEN, TET | n.d. | 0.17 |
| CDC-0178 | TET | n.d. | 0.34 |
| CDC-0179 | CIP, PEN, TET | n.d. | 0.08 |
| CDC-0180 | | n.d. | 0.17 |
| CDC-0181 | CIP, PEN, TET | n.d. | 0.17 |
| CDC-0182 | TET | n.d. | 0.17 |
| CDC-0183 | CIP, PEN, TET | n.d. | 0.34 |
| CDC-0184 | CIP, PEN, TET | n.d. | 0.17 |
| CDC-0185 | CIP, PEN, TET | n.d. | 0.27 |
| CDC-0186 | CIP, PEN, TET | n.d. | 0.17 |
| CDC-0187 | CIP, PEN, TET | n.d. | 0.34 |
| CDC-0188 | CIP, PEN | n.d. | 0.17 |
| CDC-0189 | CIP, PEN, TET | n.d. | 0.13 |
| CDC-0190 | CIP, PEN, TET | n.d. | 0.17 |
| WHO F | | n.d. | 0.06 |
| WHO G | PEN, TET | n.d. | 0.13 |
| WHO K | CIP, PEN, TET | n.d. | 0.26 |
| WHO L | CIP,PEN, TET | n.d. | 0.26 |

| | | | |
|---|---|---|---|
| WHO M | CIP,PEN,TET | n.d. | 0.13 |
| WHO N | CIP, PEN, TET | n.d. | 0.26 |
| WHO O | PEN, TET | n.d. | 0.13 |
| WHO P | PEN, TET | n.d. | 0.33 |
| WHO U | PEN, TET | n.d. | 0.13 |
| WHO V | AZM, CIP, PEN, TET | n.d. | 0.26 |
| WHO W | CIP, PEN, TET | n.d. | 0.13 |
| WHO X | CIP, PEN, TET | n.d. | 0.21 |
| WHO Y | CIP, PEN, TET | n.d. | 0.21 |
| WHO Z | CIP, PEN, TET | n.d. | 0.26 |
| MMX ATCC 49226 | | 0.25 | 0.27 |
| MMX 6744 | | 0.13 | 0.27 |
| MMX 6746 | CIP | 0.13 | 0.14 |
| MMX 6752 | TET | 0.25 | 0.27 |
| MMX 6753 | | 0.06 | 0.14 |
| MMX 6757 | | 0.06 | 0.07 |
| MMX 6758 | | 0.13 | 0.27 |
| MMX 6762 | | 0.13 | 0.27 |
| MMX 6767 | | 0.06 | 0.14 |
| MMX 6771 | TET | 0.06 | 0.14 |
| MMX 6793 | CIP | 0.06 | 0.14 |
| MMX 6797 | CIP, TET | 0.06 | 0.14 |
| MMX 6803 | CIP | 0.13 | 0.27 |
| MMX 6812 | CIP | 0.13 | 0.27 |
| MMX 6818 | CIP | 0.13 | 0.27 |
| MMX 6819 | CIP | 0.13 | 0.14 |
| MMX 6879 | | 0.06 | 0.14 |
| MMX 6921 | CIP | 0.25 | 0.53 |
| MMX 6922 | CIP | 0.25 | 0.53 |
| MMX 6983 | CIP | 0.25 | 0.53 |
| MMX 6989 | | 0.13 | 0.14 |
| MMX 6990 | CIP | 0.06 | 0.14 |
| MMX 6992 | | 0.13 | 0.14 |
| MMX 6996 | | 0.13 | 0.27 |
| MMX 6998 | CIP | 0.25 | 0.53 |
| MMX 7002 | CIP | 0.13 | 0.27 |
| MMX 7005 | | 0.03 | 0.03 |
| **$MIC_{90}$ (n)** | | **0.25 (32)** | **0.53 (71)** |
| **MIC Range (n)** | | **0.03-0.25 (32)** | **0.03-0.68 (71)** |

SPT, spectinomycin; STR, streptomycin; TET, tetracycline; PEN, penicillin G; CFM, cefixime; CRO, ceftriazone; CIP, ciprofloxacin; LVX, levofloxacin; AZM, azithromycin; MMX, Micromyx, LLC strain (these MIC assays were performed, using the broth dilution method, by Micromyx, LLC.)

**Table S4.** Antibacterial spectrum

| | | | | MIC (µg/ml) | |
|---|---|---|---|---|---|
| Category | Organism | Strain | Resistance/Description | KKL-35 | MBX-4132 |
| Gram-negative | *Escherichia coli* | KLE700 | | ≥32 | ≥35 |
| | *Escherichia coli* | KLE701 | KLE700 *ΔtolC::tet* | 0.5 | 2.7 |
| | *Klebsiella pneumoniae* | ATCC 13883 | | n.d. | ≥35 |
| | *Acinetobacter baumannii* | ATCC 19606 | | >32 | ≥35 |
| | *Pseudomonas aeruginosa* | ATCC 27853 | | >32 | ≥35 |
| | *Moraxella catarrhalis* | 8716 | | n.d. | 0.04 |
| | *Legionella pneumophila* | ATCC 33153 | | n.d. | 8.7 |
| | *Haemophilus influenzae* | ATCC 35056 | | n.d. | 17.5 |
| Gram-positive | *Staphylococcus aureus* | BAA-1717 | MRSA | n.d. | 1.21 |
| | *Staphylococcus aureus* | MRSA-1234547263 | MRSA | 1 | 3.2 |
| | *Staphylococcus aureus* | MRSA-1094 | MRSA | 1 | 3.2 |
| | *Staphylococcus aureus* | ATCC 35556 | | 1 | 4.2 |
| | *Staphylococcus aureus* | NRS-77 | | 1 | 3.2 |
| | *Staphylococcus aureus* | MSSA | | 1 | 4.2 |
| | *Staphylococcus aureus* | N315, NRS-70 | MRSA | 1.5 | 6.3 |
| | *Staphylococcus aureus* | ATCC 25923 | | 1.5 | 6.3 |
| | *Staphylococcus aureus* | THC1516 | MRSA | 0.9 | 1 |
| | *Streptococcus pneumoniae* | ATCC 49619 | | n.d. | 34.9 |
| | *Mycoplasma pneumoniae* | ATCC 15531 | | n.d. | 34.9 |

**Table S5.** Eurofins Discovery Services *In Vitro* Safety Panel, Cyp inhibition assays and Cardiac Ion Channel interaction results for MBX-4132.

| Catalog ref[A] | Receptor Profiling | Species | Conc | % inhibition |
|---|---|---|---|---|
| 104010 | Cholinesterase, Acetyl, ACES | hum | 10 µM | 6 |
| 116030 | Cycloogenase COX-1 | hum | 10 µM | 8 |
| 118030 | Cycloogenase COX-2 | hum | 10 µM | 10 |
| 140010 | Monoamine Oxidase MAO-A | hum | 10 µM | -108 |
| 152300 | Phosphodiesterase PDE3A | hum | 10 µM | -6 |
| 154420 | Phosphodiesterase PDE4D2 | hum | 10 µM | 3 |
| 176020 | Protein Tyrosine Kinase, LCK | hum | 10 µM | 29 |
| 200610 | Adenosine $A_{2A}$ | hum | 10 µM | 42 |
| 203110 | Adrenergic $\alpha_{1A}$ | hum | 10 µM | 6 |
| 203630 | Adrenergic $\alpha_{2A}$ | hum | 10 µM | 35 |
| 204010 | Adrenergic $\beta_1$ | hum | 10 µM | -8 |
| 204110 | Adrenergic $\beta_2$ | hum | 10 µM | 35 |
| 206000 | Androgen (testosterone) | hum | 10 µM | -2 |
| 214600 | Calcium Channel L-Type, Dihydropyridine | rat | 10 µM | 11 |
| 217050 | Cannabinoid $CB_1$ | hum | 10 µM | -4 |
| 217100 | Cannabinoid $CB_2$ | hum | 10 µM | -4 |
| 218030 | Cholecystokinin $CKK_1$ ($CCK_A$) | hum | 10 µM | 66 |
| 219500 | Dopamine $D_1$ | hum | 10 µM | 26 |
| 219700 | Dopamine $D_{2S}$ | hum | 10 µM | 21 |
| 224010 | Endothelin $ET_A$ | hum | 10 µM | -15 |
| 226600 | $GABA_A$, Flunitrazepam, Central | rat | 10 µM | 3 |
| 232030 | Glucocorticoid | hum | 10 µM | -2 |
| 232810 | Glutamate, NMDA, Agonism | rat | 10 µM | 8 |
| 239610 | Histamine $H_1$ | hum | 10 µM | 11 |
| 239710 | Histamine $H_2$ | hum | 10 µM | -7 |
| 252610 | Muscarinic $M_1$ | hum | 10 µM | -7 |
| 252710 | Muscarinic $M_2$ | hum | 10 µM | 7 |
| 252810 | Muscarinic $M_3$ | hum | 10 µM | 5 |
| 299031 | Nicotinic Acetylcholine $\alpha4\beta2$, Cytisine | hum | 10 µM | -5 |
| 260130 | Opiate $Delta_1$ (OP1, DOP) | hum | 10 µM | 5 |
| 260210 | Opiate Kappa (OP2, KOP) | hum | 10 µM | 13 |
| 260410 | Opiate Mu (OP3, MOP) | hum | 10 µM | 17 |

| | | | | |
|---|---|---|---|---|
| 265510 | Potassium Channel [$K_A$] | rat | 10 μM | -2 |
| 265910 | Potassium Channel hERG, [$^3$H Dofetilide | hum | 10 μM | 27 |
| 271110 | Serotonin (5-hydroxytryptamine) 5-$HT_{1A}$ | hum | 10 μM | 30 |
| 271230 | Serotonin (5-hydroxytryptamine) 5-$HT_{1B}$ | hum | 10 μM | 21 |
| 271650 | Serotonin (5-hydroxytryptamine) 5-$HT_{2A}$ | hum | 10 μM | 18 |
| 271700 | Serotonin (5-hydroxytryptamine) 5-$HT_{2B}$ | hum | 10 μM | 44 |
| 271910 | Serotonin (5-hydroxytryptamine) 5-$HT_3$ | hum | 10 μM | 4 |
| 279510 | Sodium Channel, Site 2 | rat | 10 μM | 21 |
| 220320 | Transporter, Dopamine (DAT) | hum | 10 μM | 3 |
| 204410 | Transporter, Norepinephrine (NET) | hum | 10 μM | 6 |
| 274030 | Transporter, Serotonin (5-Hyroxytryptamine) (SERT) | hum | 10 μM | -10 |
| 287530 | Vasopressin $V_{1A}$ | hum | 10 μM | -4 |

| Catalog ref[A] | Cyp Inhibition Assays | Species | $pIC_{50}$ |
|---|---|---|---|
| 4876 | CYP1A inhibition (Phenacetin substrate) | hum | NC[B] |
| 4877 | CYP2B6 inhibition (Bupropion substrate) | hum | >100 μM |
| 4879 | CYP2C8 inhibition (Amodiaquine substrate) | hum | >100 μM |
| 4878 | CYP2C9 inhibition (Diclofenac substrate) | hum | NC[B] |
| 4874 | CYP2C19 inhibition (Omeprazole substrate) | hum | >100 μM |
| 4875 | CYP2D6 inhibition (Dextromethorphan substrate) | hum | >100 μM |
| 4873 | CYP3A inhibition (Midazolam substrate) | hum | NC[B] |
| 4872 | CYP3A inhibition (Testosterone substrate) | hum | >100 μM |

| Catalog ref[A] | Cardiac Ion Channel Panel | Mode | Est. $IC_{50}$ (μM) |
|---|---|---|---|
| CYL8004QP2DR | Nav1.5 | Antagonist | >30 |
| CYL8038QP2DR | hERG | Antagonist | >30 |
| CYL8007QP2DR | KCNQ1/mink | Antagonist | >30 |
| CYL8069QP2DR | Kv4.3/ChIP2 | Antagonist | >30 (Peak&End) |
| CYL8032QP2DR | Kir2.1 | Antagonist | >30 (Peak&End) |

| | | | |
|---|---|---|---|
| CYL8051QP2DR | Cav1.2 | Antagonist | >30 |
| CYL7004QP1DR | Nav1.5 late current | Agonist | >30 |
| CYL7004QP2DR | Nav1.5 late current | Antagonist | >30 |

[A]Assays were performed using established protocols by Eurofins Discovery Services; catalog references provided. [B]No activity observed at any tested concentration.

**Table S6.** Evaluation of MBX-4132 in the *Salmonella*/Microsome Plate Incorporation Assay (AMES Screen) performed by SRI Biosciences, a division of SRI International.

| Strain | Test Compound/Condition | Dose (µg/plate) | Mean Revertants/plate |
|---|---|---|---|
| TA98 | DMSO | N/A | 23.7 ± 6.7 |
| TA100 | DMSO | N/A | 127.34.5 ± 4.5 |
| TA98 | DMSO+S9 | N/A | 28.75.9 ± 5.9 |
| TA100 | DMSO+S9 | N/A | 129.0 ± 4.4 |
| TA98 | 2-Nitrofluorene | 5 | 1331.3 ± 171.6 |
| TA100 | Sodium Azide | 5 | 2029.0 ± 49.6 |
| TA98 | 2-Aminoanthracene + S9 | 2 | 1206.7 ± 169.1 |
| TA100 | 2-Aminoanthracene + S9 | 2 | 1438.7 ± 55.8 |
| TA98 | MBX-4132 | 1 | 21.5 ± 2.1 |
| TA98 | MBX-4132 | 5 | 17.0 ± 1.4 |
| TA98 | MBX-4132 | 10 | 15.0 ± 1.4 |
| TA98 | MBX-4132 | 50 | 20.0 ± 2.8 |
| TA98 | MBX-4132 | 100 | 20.0 ± 2.8 |
| TA98 | MBX-4132 | 500 | 17.5 ± 2.1 |
| TA100 | MBX-4132 | 1 | 112.5 ± 6.4 |
| TA100 | MBX-4132 | 5 | 99.0 ± 1.4 |
| TA100 | MBX-4132 | 10 | 97.0 ± 4.2 |
| TA100 | MBX-4132 | 50 | 73.0 ± 8.5 |
| TA100 | MBX-4132 | 100 | 54.0 ± 1.4 |
| TA100 | MBX-4132 | 500 | 35.5 ± 3.5 |
| TA98 | MBX-4132 + S9 | 1 | 23.5 ± 2.1 |
| TA98 | MBX-4132 + S9 | 5 | 22.0 ± 0.0 |
| TA98 | MBX-4132 + S9 | 10 | 21.0 ± 2.8 |
| TA98 | MBX-4132 + S9 | 50 | 18.5 ± 0.7 |
| TA98 | MBX-4132 + S9 | 100 | 17.5 ± 3.5 |
| TA98 | MBX-4132 + S9 | 500 | 20.0 ± 2.8 |
| TA100 | MBX-4132 + S9 | 1 | 107.5 ± 4.9 |
| TA100 | MBX-4132 + S9 | 5 | 125.5 ± 7.8 |
| TA100 | MBX-4132 + S9 | 10 | 102.0 ± 15.6 |
| TA100 | MBX-4132 + S9 | 50 | 71.0 ± 4.2 |
| TA100 | MBX-4132 + S9 | 100 | 43.0 ± 4.2 |
| TA100 | MBX-4132 + S9 | 500 | 19.5 ± 0.7 |

**Table S7.** Results from mitchondrial toxicity studies examining the effect of MBX-4132 on reactive oxygen species (ROS), mitochondrial membrane potential (MMP) and differential cytotoxicity against HepG2 cells grown on glucose or galactose.

| Compound | Conc. (μM) | Attached live cells (%) | ROS (fold induction) | MMP inhibition (fold induction) |
|---|---|---|---|---|
| MBX-4132 | 3.18E-03 | 105.6 ± 7.5 | 3.1 ± 4.1 | 1.0 ± 0.1 |
| MBX-4132 | 1.01E-02 | 109.5 ± 8.4 | 1.4 ± 0.4 | 1.0 ± 0.1 |
| MBX-4132 | 3.18E-02 | 97.5 ± 12.2 | 0.9 ± 0.5 | 0.9 ± 0.2 |
| MBX-4132 | 1.00E-01 | 96.7 ± 6.9 | 1.4 ± 1.1 | 1.2 ± 0.3 |
| MBX-4132 | 3.17E-01 | 114.0 ± 14.4 | 1.2 ± 0.5 | 0.9 ± 0.1 |
| MBX-4132 | 1.00E+00 | 106.3 ± 3.3 | 1.2 ± 0.4 | 0.9 ± 0.1 |
| MBX-4132 | 3.17E+00 | 99.6 ± 6.1 | 1.1 ± 0.3 | 1.0 ± 0.2 |
| MBX-4132 | 1.00E+01 | 97.7 ± 9.4 | 1.9 ± 1.0 | 1.1 ± 0.1 |
| MBX-4132 | 3.16E+01 | 89.5 ± 3.6 | 1.3 ± 0.6 | 1.1 ± 0.1 |
| MBX-4132 | 1.00E+02 | 69.6 ± 12.3 | 9.2 ± 3.6 | 0.8 ± 0.1 |

| Compound | Conc. (μM) | Cell type | Medium | % viability |
|---|---|---|---|---|
| MBX-4132 | 3.00E-06 | HepG2 | Glucose | 109.6 ± 0.2 |
| MBX-4132 | 1.00E-05 | HepG2 | Glucose | 107.8 ± 2.1 |
| MBX-4132 | 3.00E-05 | HepG2 | Glucose | 106.5 ± 3.3 |
| MBX-4132 | 1.00E-04 | HepG2 | Glucose | 85.6 ± 1.8 |
| MBX-4132 | 3.00E-06 | HepG2 | Glucose | 94.1 ± 9.3 |
| MBX-4132 | 1.00E-05 | HepG2 | Glucose | 95.4 ± 0.7 |
| MBX-4132 | 3.00E-05 | HepG2 | Glucose | 96.8 ± 0.4 |
| MBX-4132 | 1.00E-04 | HepG2 | Glucose | 64.0 ± 10.3 |

**Table S8.** Data collection, model building and refinement.

| **Data collection** | **70S-P-tRNA-KKL-2098** |
|---|---|
| Voltage (keV) | 300 |
| Magnification | 59,000 |
| Electron dose ($e^-/Å^2$) | 58 |
| Pixel size (Å/pix) | 1 |
| Detector | DE64 counting mode |
| Defocus range (µm) | 1.5-3.5 |
| Micrographs | 2,197 |
| Total Particles | 474,382 |
| **Reconstruction** | |
| Particles included | 28,121 |
| $FSC_{0.5}$ | 3.8 |
| $FSC_{0.143}$ | 3.1 |
| **Model Refinement** | |
| **CCmap_model** | 0.88 |
| **Model quality** | |
| RMSD | |
| Bond lengths (Å) / Bond angles (º) | 0.007/0.897 |
| Ramachandran plot statistics | |
| Most favored (%) | 94.7 |
| Allowed | 4.22 |
| Outliers (%) | 1.12 |
| Rotamer outliers (%) | 0.3 |
| Cß outliers | 0.0 |
| Clashscore | 6.66 |

**Table S9.** Bacterial strains, plasmids, and synthetic sequences.

| strain name | description | source |
|---|---|---|
| *E. coli* BL21 (DE3) pET28-H10arfArnc | Strain for expressing non-stop ribosomes | this work |
| IW312 | *E. coli* LG90 $\Delta$*rpmA* | (*47*) |
| IW312 pL27 | contains plasmid for inducible expression of wild-type L27 | (*47*) |
| IW312 pL27 –3 | contains plasmid for inducible expression of L27 –3 | (*47*) |
| IW312 pL27 –6 | contains plasmid for inducible expression of L27 –6 | (*47*) |
| IW312 $\Delta$*tolC* pL27 | IW312 pL27 with *tolC* deleted | this work |
| IW312 $\Delta$*tolC* pL27 –3 | IW312 pL27 –3 with *tolC* deleted | this work |
| IW312 $\Delta$*tolC* pL27 –6 | IW312 pL27 –6 with *tolC* deleted | this work |
| *N. gonorrhoeae* H041 | Clinical multiple-antibiotic resistant isolate | (*10*) |
| *N. gonorrhoeae* | ATCC 49226 | ATCC |
| *N. gonorrhoeae* | ATCC 700719 | ATCC |
| *N. gonorrhoeae* | ATCC 700825 | ATCC |
| *N. gonorrhoeae* | BAA-1846 | ATCC |
| *N. gonorrhoeae* | CDC-0165 | CDC-ARBank |
| *N. gonorrhoeae* | CDC-0166 | CDC-ARBank |
| *N. gonorrhoeae* | CDC-0167 | CDC-ARBank |
| *N. gonorrhoeae* | CDC-0169 | CDC-ARBank |
| *N. gonorrhoeae* | CDC-0170 | CDC-ARBank |
| *N. gonorrhoeae* | CDC-0171 | CDC-ARBank |
| *N. gonorrhoeae* | CDC-0172 | CDC-ARBank |
| *N. gonorrhoeae* | CDC-0173 | CDC-ARBank |
| *N. gonorrhoeae* | CDC-0174 | CDC-ARBank |
| *N. gonorrhoeae* | CDC-0175 | CDC-ARBank |
| *N. gonorrhoeae* | CDC-0176 | CDC-ARBank |
| *N. gonorrhoeae* | CDC-0177 | CDC-ARBank |
| *N. gonorrhoeae* | CDC-0178 | CDC-ARBank |
| *N. gonorrhoeae* | CDC-0179 | CDC-ARBank |
| *N. gonorrhoeae* | CDC-0180 | CDC-ARBank |
| *N. gonorrhoeae* | CDC-0181 | CDC-ARBank |
| *N. gonorrhoeae* | CDC-0182 | CDC-ARBank |
| *N. gonorrhoeae* | CDC-0183 | CDC-ARBank |
| *N. gonorrhoeae* | CDC-0184 | CDC-ARBank |
| *N. gonorrhoeae* | CDC-0185 | CDC-ARBank |
| *N. gonorrhoeae* | CDC-0186 | CDC-ARBank |
| *N. gonorrhoeae* | CDC-0187 | CDC-ARBank |
| *N. gonorrhoeae* | CDC-0188 | CDC-ARBank |
| *N. gonorrhoeae* | CDC-0189 | CDC-ARBank |
| *N. gonorrhoeae* | CDC-0190 | CDC-ARBank |
| *N. gonorrhoeae* | WHO F | CDC-ARBank |
| *N. gonorrhoeae* | WHO G | CDC-ARBank |
| *N. gonorrhoeae* | WHO K | CDC-ARBank |

| | | |
|---|---|---|
| *N. gonorrhoeae* | WHO L | CDC-ARBank |
| *N. gonorrhoeae* | WHO M | CDC-ARBank |
| *N. gonorrhoeae* | WHO N | CDC-ARBank |
| *N. gonorrhoeae* | WHO O | CDC-ARBank |
| *N. gonorrhoeae* | WHO P | CDC-ARBank |
| *N. gonorrhoeae* | WHO U | CDC-ARBank |
| *N. gonorrhoeae* | WHO V | CDC-ARBank |
| *N. gonorrhoeae* | WHO W | CDC-ARBank |
| *N. gonorrhoeae* | WHO X | CDC-ARBank |
| *N. gonorrhoeae* | WHO Y | CDC-ARBank |
| *N. gonorrhoeae* | WHO Z | CDC-ARBank |
| *N. gonorrhoeae* | MMX 6744 | Micromyx, LLC |
| *N. gonorrhoeae* | MMX 6746 | Micromyx, LLC |
| *N. gonorrhoeae* | MMX 6752 | Micromyx, LLC |
| *N. gonorrhoeae* | MMX 6753 | Micromyx, LLC |
| *N. gonorrhoeae* | MMX 6757 | Micromyx, LLC |
| *N. gonorrhoeae* | MMX 6758 | Micromyx, LLC |
| *N. gonorrhoeae* | MMX 6762 | Micromyx, LLC |
| *N. gonorrhoeae* | MMX 6767 | Micromyx, LLC |
| *N. gonorrhoeae* | MMX 6771 | Micromyx, LLC |
| *N. gonorrhoeae* | MMX 6793 | Micromyx, LLC |
| *N. gonorrhoeae* | MMX 6797 | Micromyx, LLC |
| *N. gonorrhoeae* | MMX 6803 | Micromyx, LLC |
| *N. gonorrhoeae* | MMX 6812 | Micromyx, LLC |
| *N. gonorrhoeae* | MMX 6818 | Micromyx, LLC |
| *N. gonorrhoeae* | MMX 6819 | Micromyx, LLC |
| *N. gonorrhoeae* | MMX 6879 | Micromyx, LLC |
| *N. gonorrhoeae* | MMX 6921 | Micromyx, LLC |
| *N. gonorrhoeae* | MMX 6922 | Micromyx, LLC |
| *N. gonorrhoeae* | MMX 6983 | Micromyx, LLC |
| *N. gonorrhoeae* | MMX 6989 | Micromyx, LLC |
| *N. gonorrhoeae* | MMX 6990 | Micromyx, LLC |
| *N. gonorrhoeae* | MMX 6992 | Micromyx, LLC |
| *N. gonorrhoeae* | MMX 6996 | Micromyx, LLC |
| *N. gonorrhoeae* | MMX 6998 | Micromyx, LLC |
| *N. gonorrhoeae* | MMX 7002 | Micromyx, LLC |
| *N. gonorrhoeae* | MMX 7005 | Micromyx, LLC |
| *Escherichia coli* | KLE700 | (*48*) |
| *Escherichia coli* | KLE701 *tolC::tet* | (*48*) |
| *Klebsiella pneumoniae* | ATCC 13883 | ATCC |
| *Acinetobacter baumannii* | ATCC 19606 | ATCC |
| *Pseudomonas aeruginosa* | ATCC 27853 | ATCC |
| *Moraxella catarrhalis* | ATCC 8716 | ATCC |
| *Legionella pneumophila* | ATCC 33153 | ATCC |
| *Haemophilus influenzae* | ATCC 35056 | ATCC |

| | | | |
|---|---|---|---|
| *Staphylococcus aureus* | BAA-1717 | | ATCC |
| *Staphylococcus aureus* | MRSA-1234547263 | | (*49*) |
| *Staphylococcus aureus* | MRSA-1094 | | (*50*) |
| *Staphylococcus aureus* | ATCC 35556 | | ATCC |
| *Staphylococcus aureus* | NRS-77 | | BEI Resources |
| *Staphylococcus aureus* | MSSA | | (*50*) |
| *Staphylococcus aureus* | N315, NRS-70 | | BEI Resources |
| *Staphylococcus aureus* | ATCC 25923 | | ATCC |
| *Staphylococcus aureus* | THC1516 | | (*51*) |
| *Streptococcus pneumoniae* | ATCC 49619 | | ATCC |
| *Mycoplasma pneumoniae* | ATCC 15531 | | ATCC |
| plasmid name | description | | source |
| pET28-H10arfArnc | plasmid to produce non-stop ribosomes *in vivo* | | this work |
| pNL3.1 | nano-luciferase encoding plasmid | | Promega |
| pMC1 | Nano-luciferase gene cloned into the NcoI and BamH1 sites of pET28 | | this work |
| DNA name | description | sequence | source |
| T7 universal | primer | TAATACGACTCACTATAGGG | ThermoFisher |
| nanoluc-ns | primer | CCCCCCGGTTACCCGGAAGAGCAGGGAGCCGTC | this work |
| nanoluc-stop | primer | TTACAGAATCTCCTCGAACAGCCG | this work |
| tmRNA-nl template | synthetic DNA cassette | GGGGCTGATTCTGGATTCGACGGGATTTGCGAAACCCAAGGTGCATGCCGAGGGGCGGTTGGCCTCGTAAAAAGCCGCAAAAAATAGTCGCAGTCTCCGGATGGCGCCTTTTTAAAAAAATTTCTTAATAACAATTTTTTTAGCCCTCTCTCCCTAGCCTCCGCTCTTAGGACGGGGATCAAGAGAGGTCAAACCCAAAAGAGATCGCGTGGAAGCCCTGCCTGGGGTTGAAGCGTTAAAACTTAATCAGGCTAGTTTGTTAGTGGCGTGTCCGTCCGCAGCTGGCAAGCGAATGTAAAGACTGACTAAGCATGTAGTACCGAGGATGTAGGAATTTCGGACGCGGGTTCAACTCCCGCCAGCTCCACCA | this work |

### SUPPLEMENTAL FIGURES

**A.**

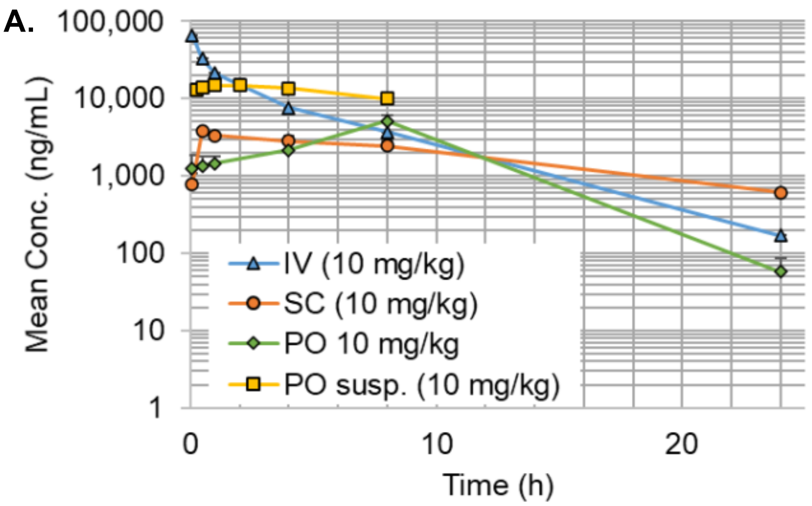

**B.**

| | $T_{1/2}$ (hr) | $C_{0/max}$ (ng/mL) | $AUC_{last}$ (hr*ng/mL) | $V_{ss}$ (mL/kg) | Cl (mL/hr/kg) | %F (Last) |
|---|---|---|---|---|---|---|
| 10 mg/kg IV | 3.55 | 76,507 | 137,462 | 266 | 72.6 | - |
| 10 mg/kg SC | 8.76 | 3,793 | 46,916 | - | - | 34 |
| 25 mg/kg SC | - | 5,100 | 76,085 | - | - | - |
| 10 mg/kg PO | - | 5,110 | 62,617 | - | - | 46 |
| 10 mg/kg susp. PO | - | 14,812 | 181,695 | - | - | >95* |

**Supplemental Figure S1. Pharmacokinetic properties of MXB-4132** (A) Graphical presentation of murine plasma concentration over time for MBX-4132. For clarity, error bars are only shown in the positive direction. “susp.” Refers to the suspension formulation studies performed at Neosome, all other data is from liquid formulation performed at Charles River Labs. (B) Calculated parameters for each dosing regimen of MBX-4132. %F for the 10 mg/kg PO suspension formulation is an estimate based on the IV data for different mouse species and gender.

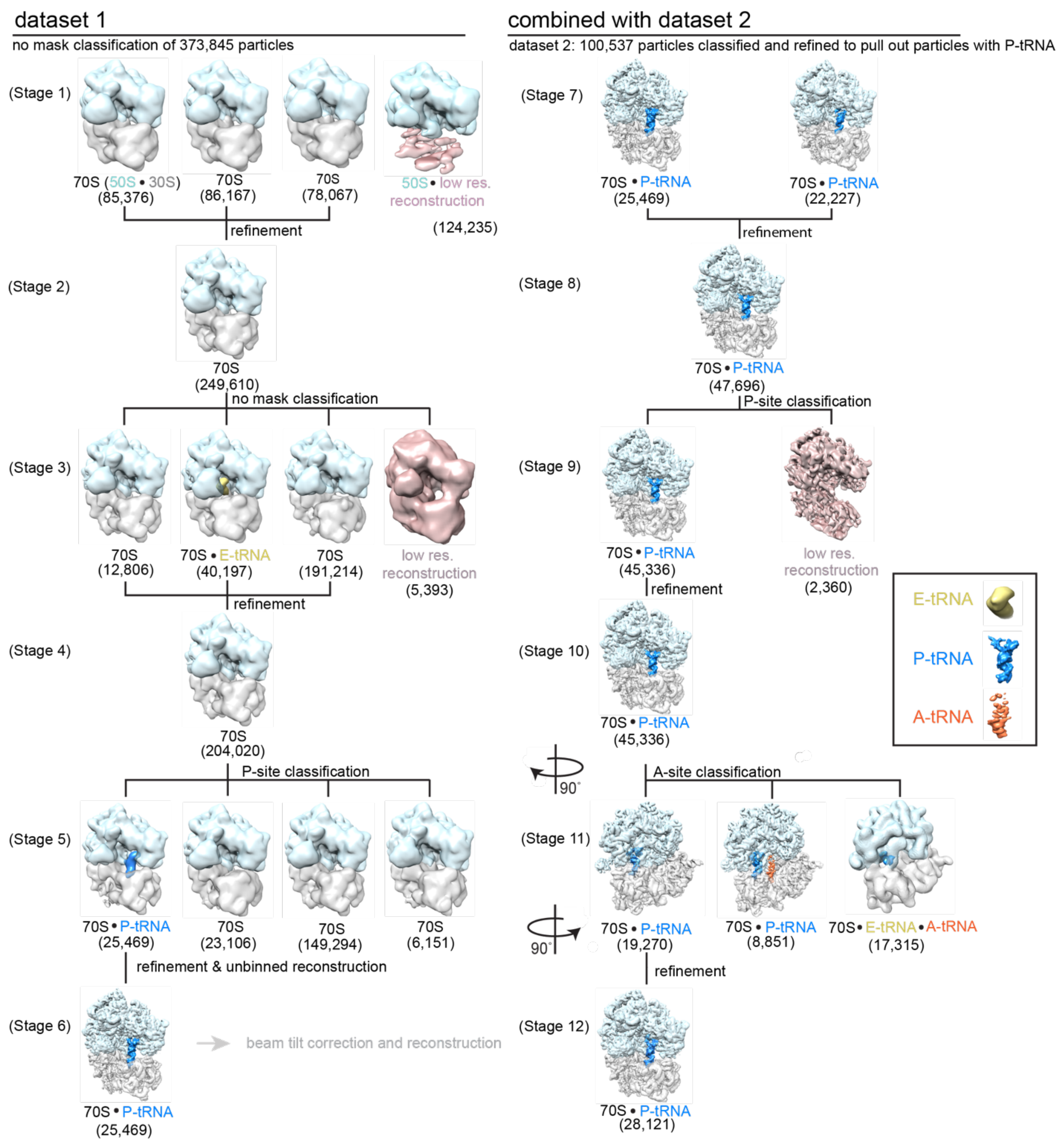

**Supplemental Figure S2. Classification of cryo-EM datasets of *E. coli* 70S complex containing P-site tRNA and KKL-2098**. To extract 70S particles with P-site tRNA and the KKL-2098 molecule several classification and refinement steps were performed. Stage 1) All particles were refined and reconstructed, then classification without alignment was performed to split the particles into 4 classes. Stage 2) 50S particles from stage 1 (shown in blue) were removed and the remaining particles refined. Stage 3) The aligned particles from stage 2 were classified into 4 classes. Stage 4) Particles contributing to the low-resolution reconstruction (pink) and 70S structures containing E-site tRNA (yellow) were removed and the remaining particles refined. Stage 5) Particles were classified without alignment into 4 classes using a P-site mask to identify particles with P-tRNA (blue). Stage 6) Particles containing P-tRNA were combined, unbinned and refined. Stage 7) Particles were beam-tilt corrected. Stage 8) Particles from both datasets were combined and refined together. Stage 9) The combined particles were classified using a P-site mask. Stage 10) Particles contributing to the lower-resolution reconstruction were eliminated and 70S/P-tRNA particles were refined. Stage 11) The reconstruction from stage 10 contained some A-site tRNA density, so a focused classification without alignment using an A-site mask was performed. Ribosomes are shown at 90° rotated view in respect to stage 10. Stage 12) 70S particles containing A-site tRNA (orange) were eliminated and 70S/P-tRNA particles were refined to an overall resolution of 3.2 Å. The numbers of particles that make up each reconstruction are depicted for each complex.

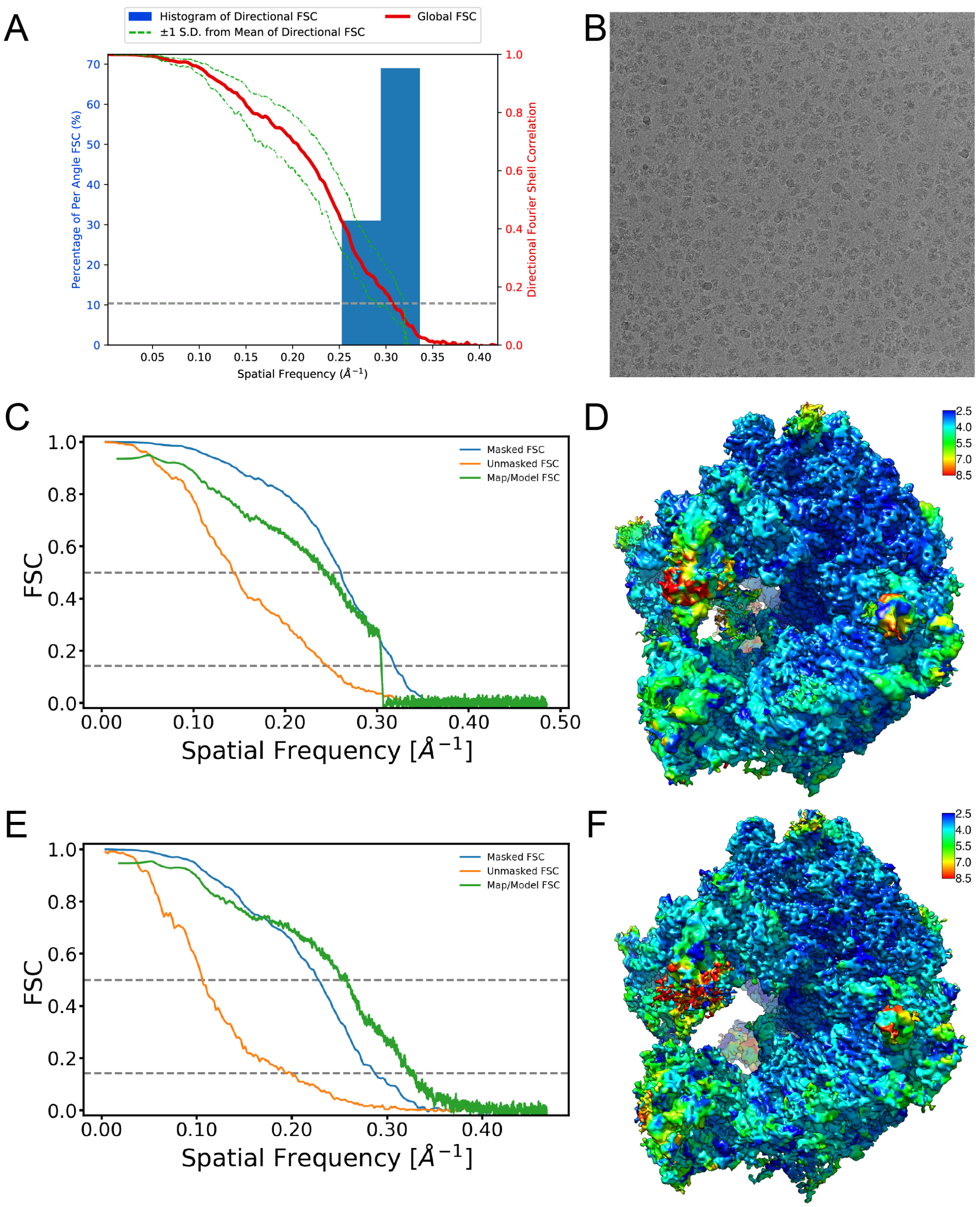

**Supplemental Figure S3. Resolution characterization and quality of cryo-EM maps.** (A) 3D FSC plot (*14*) showing degree of directional anisotropy for the 70S•P-tRNA•KKL map. (B) A representative micrograph from the cryo-EM dataset. (C) FSC plots for the 70S•P-tRNA•KKL map for the independent half maps (orange) masked (cyan), and map/model FSC (green). Dotted lines are shown at $FSC_{0.5}$ and $FSC_{0.143}$. (D) Local resolution estimate for the 70S-P-tRNA-KKL-2098 map estimated from blocres (*15*) . The map is colored from highest resolution (blue) to lowest resolution (red). (E) FSC plots for the 70S-KKL-2098 map for the independent half maps (orange) masked (cyan), and map/model FSC (green). (F) Local resolution estimate for the 70S•KKL map colored as in D.

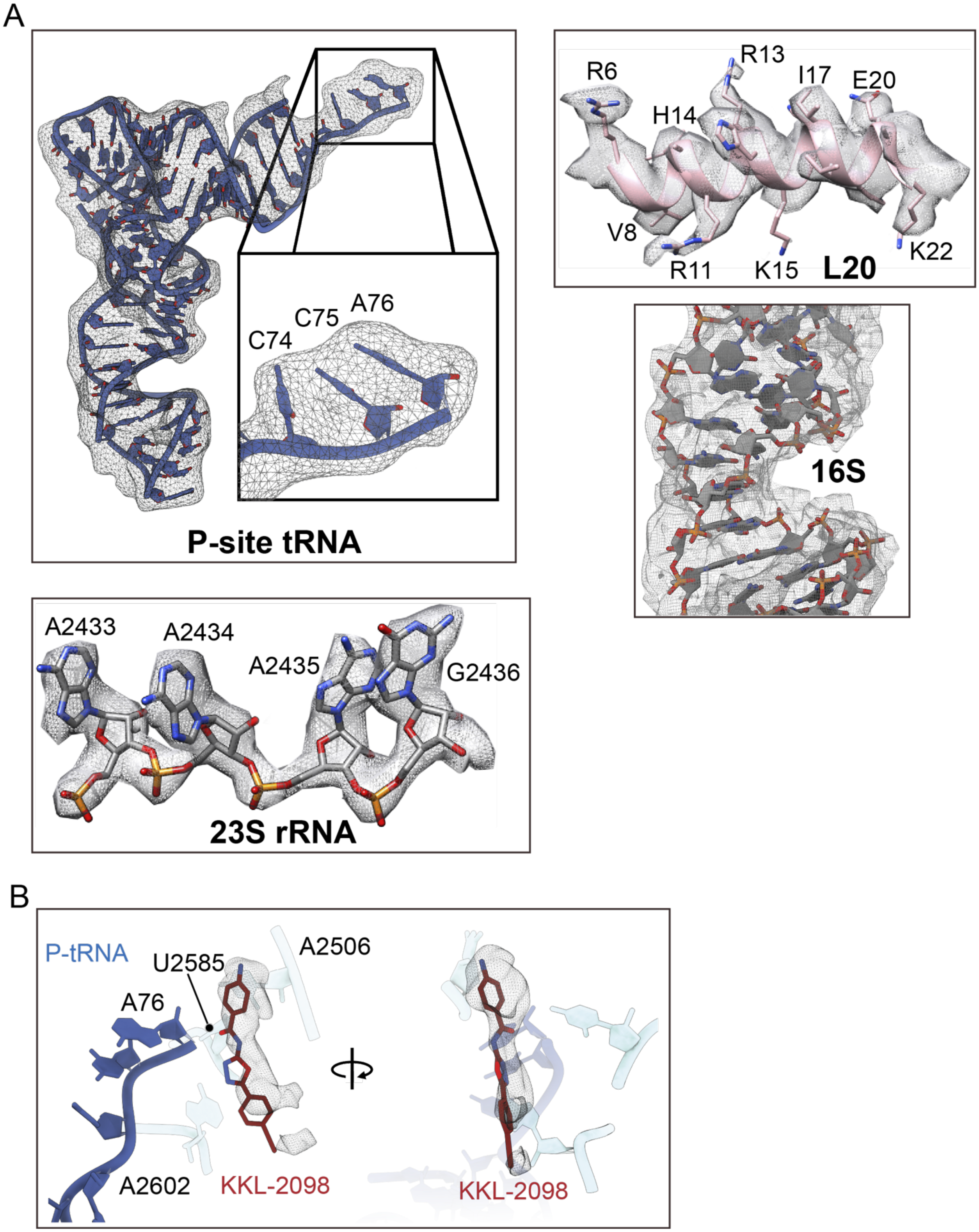

**Supplemental Figure S4. Data quality of representative areas of the 70S-P-tRNA-KKL-2098 map.** (A) Representative map quality of the P-site tRNA with the inset showing the CCA end. Map quality for ribosomal protein L20, 16S rRNA and 23S rRNA. (B) Map quality of KKL-2098.

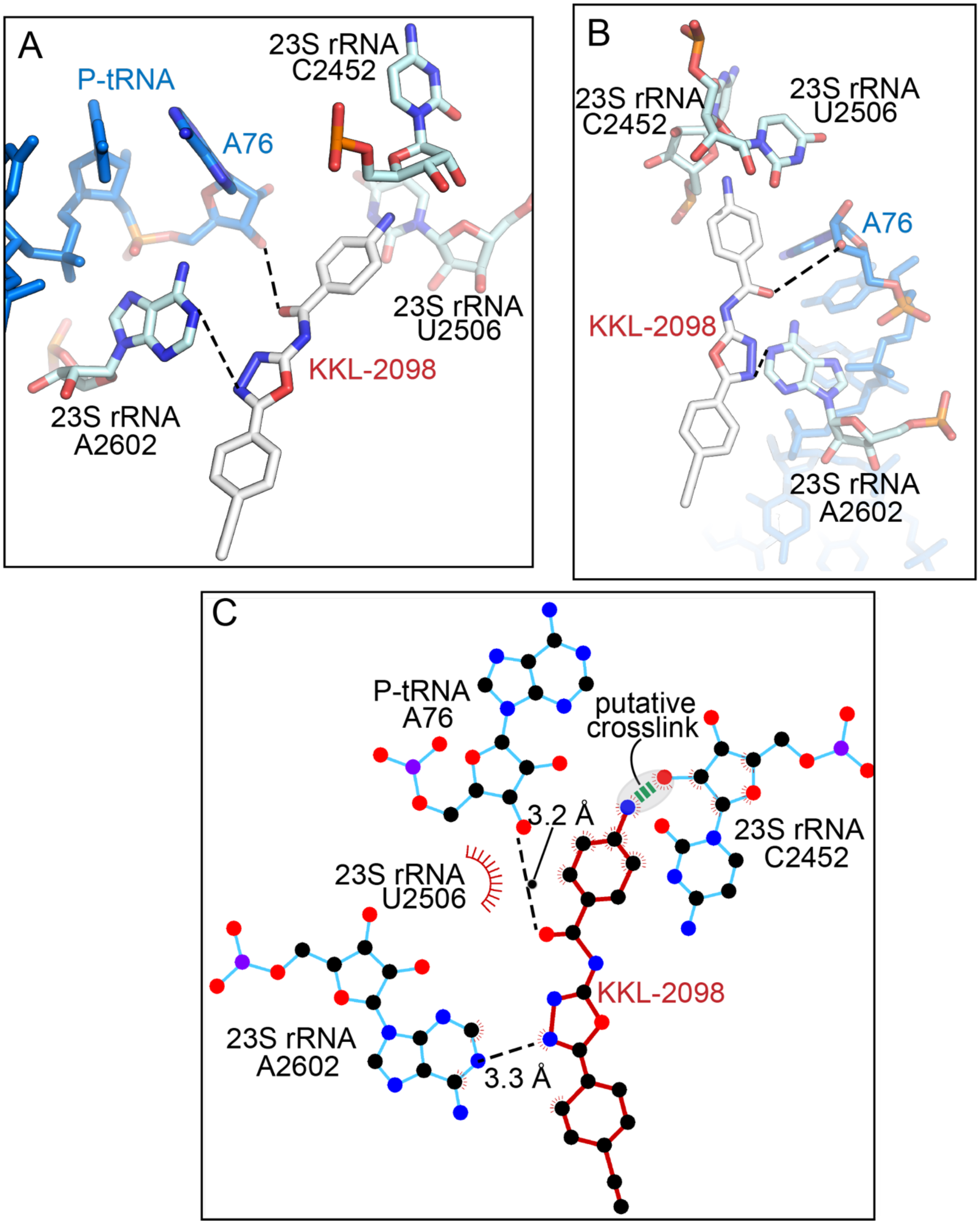

**Supplemental Figure S5. Interaction network of KKL-2098 with the ribosome. (**A) Interactions with of KKL-2098 with P-site tRNA A76 and 23 rRNA nucleotides C2452, A2506, and A2602 with a ~90º rotation in panel B. (C) A 2-dimensional representation of these interactions using LigPlot+ (*52*).

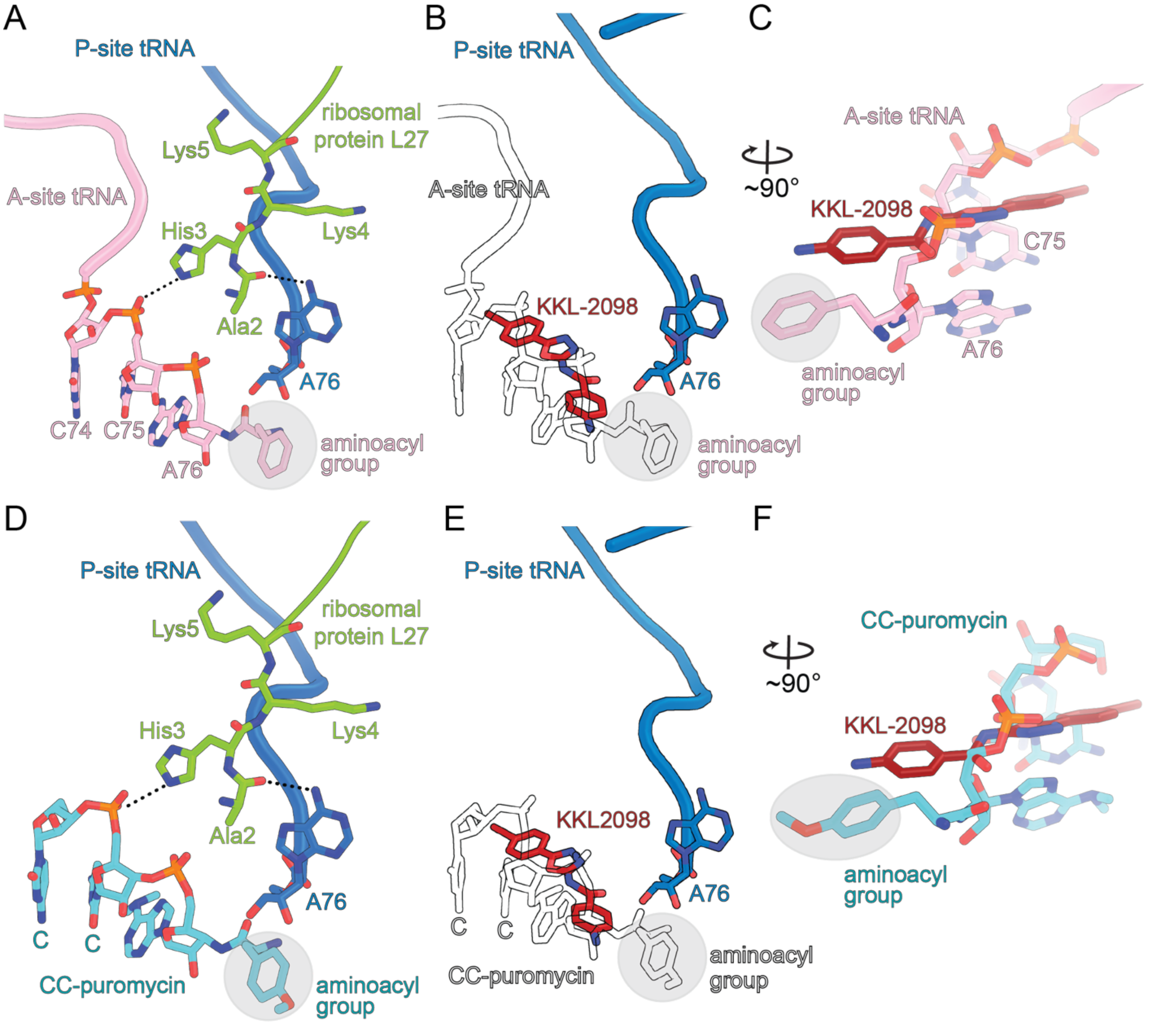

**Supplemental Figure S6. Comparison of KKL-2098 to A-site tRNA and CC-puromycin locations.** (A) The N terminus of L27 is stabilized when an aminoacylated (aa)-tRNA is bound at the A site (PDB ID 4V5C). (B) Overlay of aa-tRNA and KKL-2098. (C) A ~90º rotation of panel B shows how KKL-2098 overlaps primarily with the phosphate of A76. (D) The stabilization of the N terminus of L27 also occurs when CC-puromycin is bound (PDB ID 6OTR). (E) Overlay of CC-puromycin and KKL-2098. (F) A ~90º rotation of panel B shows how KKL-2098 overlaps primarily with the phosphate of CC-puromycin.

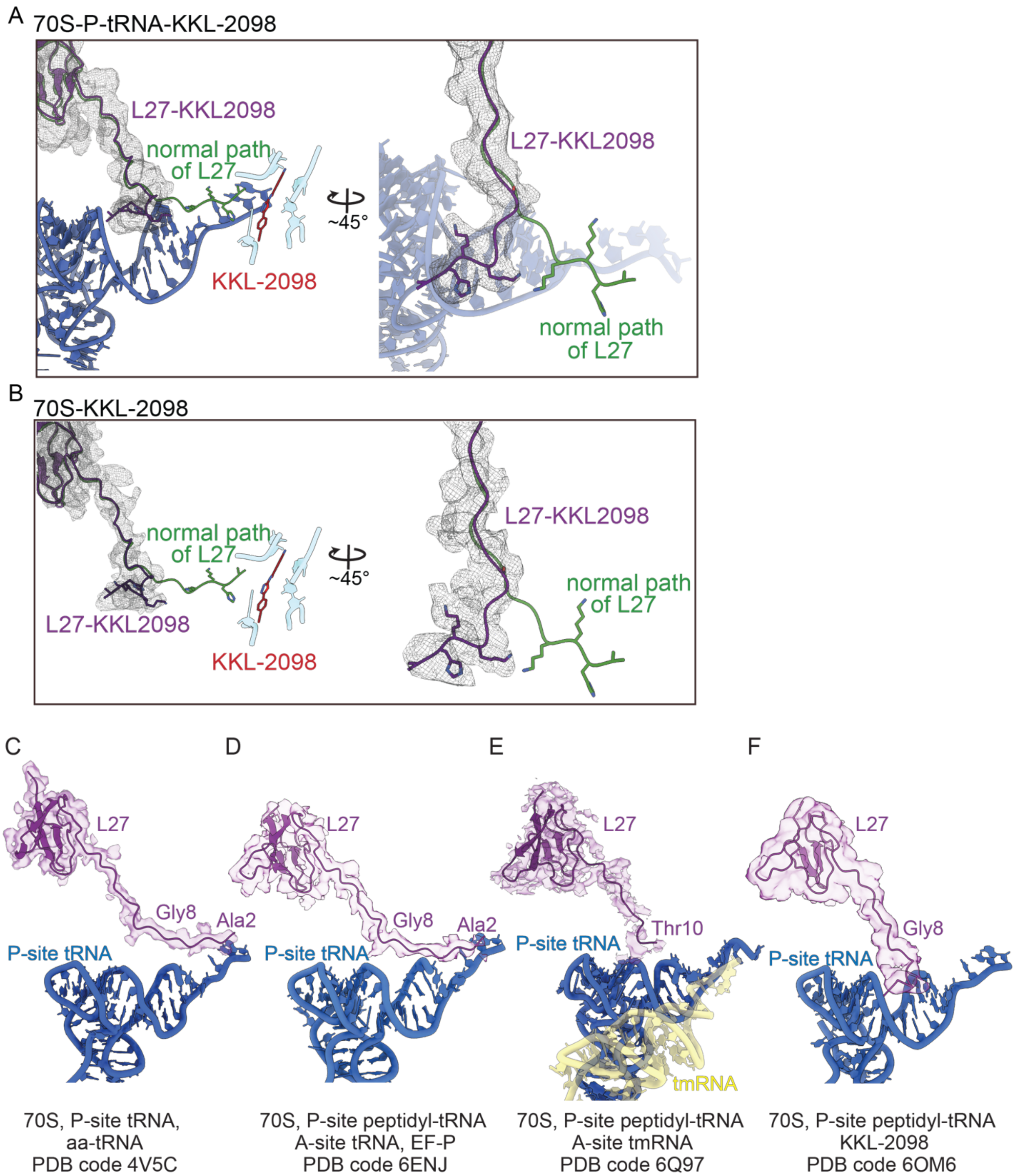

**Supplemental Figure S7. L27 map quality and its location in other structures.** (A) In the 70S-P-site tRNA-KKL-2098 structure, the N terminus of L27 (purple) moves ~180º away from the PTC. The previously observed position of L27 is shown in green (PDB ID 6ENU). (B) In the 70S-KKL-2098 structure, L27 adopts a similar position as when P-site tRNA-KKL-2098 is bound as shown in panel A. It is proposed that these particles represent a complex where peptidyl-P-site tRNA has dropped off during preparation. Position and electron potential maps of L27 in PDB code 4V5C (C), 6ENJ (D), 6Q97 (E) and 6OM6 (F).

6. Clinical and Laboratory Standards Institute, Ed., *Methods for dilution antimicrobial susceptibility tests for bacteria that grow aerobically: M07-A10 ; approved standard* (Committee for Clinical Laboratory Standards, Wayne, PA, 10. ed., 2015), *Documents / Clinical and Laboratory Standards Institute*.

12. A. S. Dixon, M. K. Schwinn, M. P. Hall, K. Zimmerman, P. Otto, T. H. Lubben, B. L. Butler, B. F. Binkowski, T. Machleidt, T. A. Kirkland, M. G. Wood, C. T. Eggers, L. P. Encell, K. V. Wood, NanoLuc

Complementation Reporter Optimized for Accurate Measurement of Protein Interactions in Cells. *ACS Chem. Biol.* **11**, 400–408 (2016).

13. T.-W. Kim, J.-W. Keum, I.-S. Oh, C.-Y. Choi, C.-G. Park, D.-M. Kim, Simple procedures for the construction of a robust and cost-effective cell-free protein synthesis system. *J. Biotechnol.* **126**, 554–561 (2006).

14. N. S. Ramadoss, J. N. Alumasa, L. Cheng, Y. Wang, S. Li, B. S. Chambers, H. Chang, A. K. Chatterjee, A. Brinker, I. H. Engels, K. C. Keiler, Small molecule inhibitors of trans-translation have broad-spectrum antibiotic activity. *Proc. Natl. Acad. Sci. U. S. A.* **110**, 10282–10287 (2013).

15. C. J. Woolstenhulme, S. Parajuli, D. W. Healey, D. P. Valverde, E. N. Petersen, A. L. Starosta, N. R. Guydosh, W. E. Johnson, D. N. Wilson, A. R. Buskirk, Nascent peptides that block protein synthesis in bacteria. *Proc. Natl. Acad. Sci.* **110**, E878–E887 (2013).

16. Clinical and Laboratory Standards Institute, *Performance standards for antimicrobial susceptibility testing* (2017).
